# Multiomic analysis of clonal development reveals new regulators of leukemic cell growth

**DOI:** 10.1101/2024.02.29.582846

**Authors:** Gracia Bonilla, Alexander Morris, Sharmistha Kundu, Anthony DuCasse, Grace Kirkpatrick, Nathan E. Jeffries, Kashish Chetal, Emma E. Yvanovich, Jelena Milosevic, Ting Zhao, Jun Xia, Rana Barghout, David Scadden, Michael K. Mansour, Robert E. Kingston, David B. Sykes, Francois E. Mercier, Ruslan I. Sadreyev

## Abstract

Mechanisms driving the increase of cell growth in developing leukemia are not fully understood. We focused on epigenomic regulation of this process by analyzing the changes of chromatin marks and gene expression in leukemic cell clones as they progressed towards increased proliferation in a mouse model of acute myeloid leukemia (AML). This progression was characterized by gradual modulation of chromatin states and gene expression across the genome, with a surprising preferential trend of reversing the prior changes associated with the origins of leukemia. Our analyses of this modulation in independently developing clones predicted a small set of potential growth regulators whose transcriptomic and epigenomic progression was consistent between clones and maintained both in vivo and ex vivo. We selected three of these genes as candidates (Irx5 and Plag1 as growth suppressors and Smad1 as a driver) and successfully validated their causal growth effects by overexpression in leukemic cells. Public patient data confirmed expression levels of IRX5 and SMAD1 as markers of AML status and survival, suggesting that multiomic analysis of evolving clones in a mouse model is a valuable predictive approach relevant to human AML.

## INTRODUCTION

Challenges in treating acute myeloid leukemia (AML) often arise from disease heterogeneity and clonal evolution during disease development [1,2] and post-treatment relapse [3,4]. This development of leukemic cell clones towards higher proliferation is driven by both genetic and epigenetic changes in hematopoietic stem cells and leukemic progenitors [1,5]. Epigenetic dysregulation is a major component of AML origin and development. Genome-wide dynamics of DNA methylation [6] linked to methylases [7] and demethylase proteins [8] are associated with AML progression and outcome. COMPASS and SWI-SNF chromatin modifiers[9] that include MLL and other protein components [10,11], COMPASS-associated Menin-LEDGF complex [12], and COMPASS antagonists, Polycomb repressive complexes [13–15] are involved both in the origin and further development of AML. Another COMPASS antagonist, histone demethylase Kdm5b is a strong suppressor of AML aggressiveness[15]. Despite clinical and functional evidence of a central role that DNA and chromatin modifiers play in the evolution and maintenance of leukemia, understanding of epigenetic evolution among individual AML clones has remained elusive, in large part due to the heterogeneity of both molecular lesions and phenotypes across individual subclones [1,2,5,16,17].

Addressing these limitations, we analyzed a tractable clonal model to understand the dynamics of chromatin regulation and gene expression during clonal evolution and to identify candidate drivers and suppressors of this process. We used a well-established mouse model of AML driven by the MLL-AF9 fusion [18], a prevalent MLL rearrangement observed in AML patients [19]. In this basic cellular model, we tracked the evolution of individual leukemic clones as they competed in mouse bone marrow, in order to identify causal regulators of clonal growth. We assumed that epigenomic regulation is a major driving process in clonal evolution and therefore performed genome-wide profiling of chromatin marks and gene expression as these clones evolved towards higher proliferation rates. Our integrative analysis of these multiomics data revealed gradual shifts of expression levels and chromatin states during clonal evolution. Surprisingly, most of these shifts were reversing the prior changes observed at the early leukemic stage.

We further leveraged the heterogeneity of clonal evolutionary paths to identify a small set of core genes whose expression and chromatin changes were consistent between individual clones. We hypothesized that these core genes were enriched in shared regulators of clonal growth. We selected and validated three of these genes, Irx5 and Plag1 as cell growth suppressors and Smad1 as a driver. The downstream transcriptional effects of these three genes converged on a group of major factors that may represent a broader regulatory network of clonal growth. Assessment of gene expression and survival data in AML patients confirmed IRX5 and SMAD1 expression levels as markers of aggressiveness status in human leukemia. These results suggest that in-depth multiomic analyses of clonal leukemic growth in a basic mouse cell model may serve as a valuable tool to identify new markers in human AML.

## RESULTS

### Chromatin and gene expression profiling at two stages of clonal evolution

We employed a cell model of clonal evolution based on the leukemic progenitors from the mouse AML model driven by the human MLL-AF9 (KMT2A-MLLT3) fusion protein [18], a recurrent MLL rearrangement [19] which drives the reprogramming of committed granulocyte macrophage progenitors (GMPs) into leukemic stem cells (LSCs) [18,20]. To label and track individual leukemic clones, GMPs isolated from CD45.1 congenic mouse were transduced with four individual fluorescent proteins. An equal mix of labeled clones of four colors was transplanted into irradiated host mice, followed by longitudinal tracking in bone marrow and blood for 120 days (Figure 1A). Following engraftment, these clones evolved in bone marrow, resulting in cell-autonomous differences in growth rate. The faster growing clones ‘won’ the competition and dominated the bone marrow and peripheral blood.

**Figure 1.**
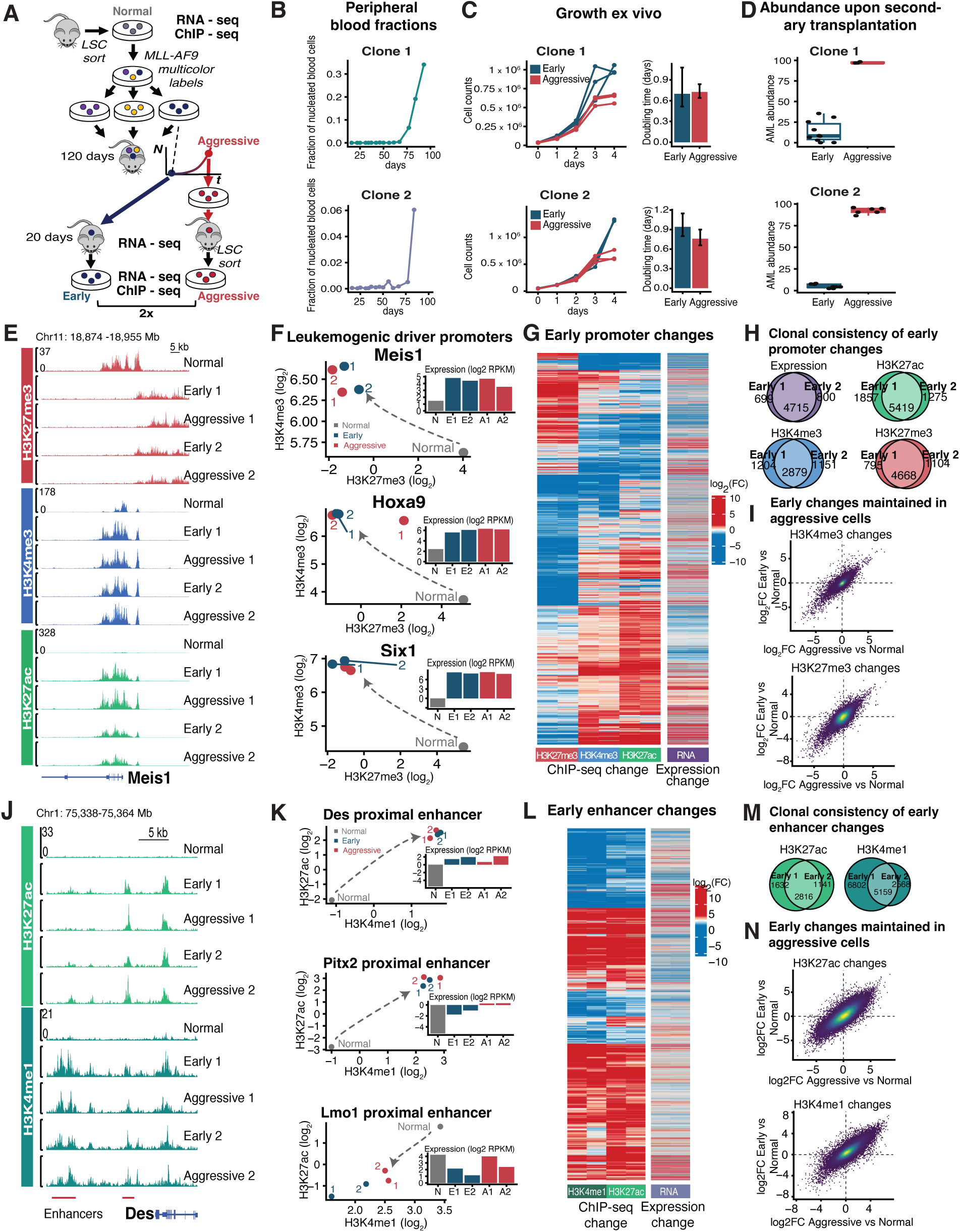
Chromatin and gene expression changes between normal and early leukemic progenitors were consistent between independent clones and mostly maintained through clonal evolution. **A.** A mix of leukemic clones individually labeled with different fluorescent proteins was transplanted in a primary mouse recipient, followed by clonal evolution for 120 days and collection of the ‘winning’ rapidly growing subclones, which maintained their aggressive growth upon secondary transplantation. The early and aggressive stages of two leukemic clones were profiled by RNA-seq in vivo and ex vivo, and by ChIP-seq of histone marks ex vivo, and compared to normal progenitors. **B-D.** Cell growth of early leukemic (blue) and aggressive (red) subclones of two individual MLLr progenitor clones (top and bottom row) during clonal selection upon primary transplantation in bone marrow (**B**), upon extraction and culturing ex vivo (**C**, doubling times as barplots), and upon secondary transplantation (**D**, abundance in bone marrow). **E.** Promoter chromatin state of a major leukemogenic driver Meis1 was dramatically activated in leukemic cells compared to normal but showed no strong difference between two stages of clonal evolution. ChIP-seq genomic tracks of H3K27me3 (red), H3K4me3 (blue), and H3K27ac (green) in normal progenitors (top) compared to two independent leukemic clones 1 and 2 at early and aggressive stages. **F.** Known drivers of MLLr leukemia were activated at the early stage but did not show further changes during clonal evolution. Promoter chromatin state progression for Meis1 (tracks in **E**), HoxA9, and Six1 (tracks in Figure S1) in the space of H3K27me3 (x axis) vs H3K4me3 density (y axis) at TSS-proximal region. Normal progenitors and early and aggressive stages of two leukemic clones (1 and 2) are represented as grey, blue, and red points, respectively. Gene expression levels (RNA-seq RPKM) are shown as barplots. Early loss of repressive H3K27me3 mark accompanied by strong gain of active H3K4me3 mark (arrows) and upregulation of expression were later maintained at the aggressive stage. **G.** Heatmap of early changes (log2 fold change) of promoter chromatin marks across all promoters with a substantive absolute level and > 2-fold change of at least one mark, juxtaposed with the changes of gene expression in vivo (log2 fold change, right). Changes were strongly consistent in two independent clones (two columns for each mark and expression). **H.** Venn diagrams of a strong overlap between the early stages of two independent clones (Early 1, Early 2) among promoters with differential gene expression in vivo (purple) and chromatin mark density (green, blue, and red) compared to normal progenitors. **I.** Most of the early changes of promoter chromatin marks and expression were later maintained at the same level during clonal evolution. Scatterplots of chromatin mark changes (log2 fold change) at all promoters between normal cells and the early leukemic stage (y axis) compared to the changes between normal cells and the aggressive leukemic stage (x axis). Individual promoters are represented as points, color indicates point density. **J.** Example of ChIP-seq tracks at two enhancers that changed chromatin state at the early leukemic stage. H3K27ac (green) and H3K4me1 (dark green) densities in normal progenitors (top) compared to two independent leukemic clones 1 and 2 at the early and aggressive stage. **K.** Chromatin state progression at enhancers proximal to Des (shown in **J**), Lmo1, and Pitx2 genes in the space of H3K4me1 (x axis) vs H3K27ac density (y axis). Points represent normal progenitors (grey) and the early (blue) and aggressive (red) stages of two leukemic clones (1 and 2). **L.** Heatmap of the early leukemic changes (log2 fold change) of H3K4me1 and H3K27ac densities across all enhancers with a substantive absolute level and > 2-fold change of at least one mark, juxtaposed with the changes of proximal gene expression in vivo (log2 fold change, right). Changes were largely consistent between marks and between two independent clones (two columns for each mark and expression). **M.** Venn diagrams of a strong overlap between two clones (Early 1, Early 2) among enhancers with chromatin mark changes between normal progenitors and the early leukemic stage. **N.** Most of the early changes of enhancer chromatin marks were later maintained during clonal evolution. Genome-wide scatterplots of chromatin mark changes (log2 fold change) at all enhancers between normal cells and the early leukemic stage (y axis) compared to the changes between normal cells and the aggressive leukemic stage (x axis). Individual enhancers are represented as points; color indicates point density.

To understand the epigenetic and transcriptional regulation during this clonal evolution, we focused on two independent leukemias that evolved into more aggressive subclones (Figures 1A,B). As they evolved from the early leukemic stage to a more aggressive stage, these winning clones dramatically increased growth rate in vivo (Figures 1A,B). Notably, this growth difference between the early and aggressive subclones was significantly diminished ex vivo (Figure 1C), suggesting the essential role of the bone marrow microenvironment in maintaining increased growth. This difference, however, was restored upon secondary re-transplantation (Figure 1D), suggesting an intrinsic cellular memory of the evolved clonal state that was maintained over multiple cell divisions ex vivo. We performed RNA-seq profiling of gene expression in LSC-enriched cell populations at the early and more aggressive stages of the same clone both in vivo and ex vivo, as well as ChIP-seq profiling of H3K4me1, H3K4me3, H3K27ac, and H3K27me3 chromatin marks ex vivo. LSC-enriched progenitors from healthy mice were profiled as a normal comparator (Figure 1A).

### Early leukemic changes of promoter chromatin marks and gene expression were consistent between independent clones

Most of the leukemia-associated changes of gene expression and promoter chromatin marks were maintained during clonal evolution. As an example of a major driver of MLL-rearranged (MLLr) leukemia [21], *Meis1* gene was strongly activated at the early leukemic stage. This was evidenced by the upregulation of *Meis1* expression compared to normal progenitors, concomitant with the loss of repressive chromatin mark H3K27me3 and gain of active marks H3K27ac and H3K4me3 at *Meis1* promoter (Figures 1E, F). However, the activation levels of *Meis1* and other major leukemic drivers did not change in the later aggressive subclones (Figures 1E,F, S1A-E), suggesting that their further upregulation is not required for clonal evolution.

Genome-wide, the changes of promoter chromatin marks and gene expression between normal progenitors and the early leukemic stage were mostly consistent between independent clones and maintained throughout leukemia progression of the same clone (Figures 1G, S1F). When compared to normal progenitors, the two early-stage leukemic clones shared the vast majority (∼85%) of differentially expressed genes (DEGs, Supplementary Data 1) and large fractions of promoters with H3K27ac, H3K4me3, and H3K27me3 changes (Figure 1H). According to EnrichR [22] pathway analysis, DEGs shared between clones were enriched in IL2/STAT5 and EMT pathways and targets of central transcription factors (Myc, Hif1a, Sox2, Tcf3, Mecom, Af4, Runx1, Cebpa, Irf8, p53, Smad3) and epigenetic factors (p300, Brd4, Polycomb proteins, Dnmt1) implicated in hematopoiesis and leukemogenesis (Figure S1G).

Compared to normal progenitors, promoter H3K27ac was preferentially increased and H3K7me3 was preferentially decreased, whereas H3K4me3 changes were more balanced (Figure S1H). Global gene expression was modestly skewed towards upregulation (Figure S1H). The combination of chromatin marks was robustly predictive of expression changes, with active marks H3K27ac and H3K4me3 being the strongest predictors (Figure S1J). However, not all promoter chromatin remodeling manifested in expression changes. For example, many transcriptionally silent promoters maintained their silent state after undergoing H3K27me3 decrease without increasing active H3K4me3 and H3K27ac marks (Figure 1G, middle part of the heatmap).

Early and aggressive subclones of the same leukemic clone were mostly similar in their genome-wide differences of expression and promoter chromatin marks compared to normal progenitors (Figure 1I, S1I, points near diagonal), suggesting that their leukemic clonal state was largely maintained through clonal evolution. However, a minority of promoters deviated between the two stages (Figure 1I, S1I, points away from diagonal) and were the focus of our further analyses.

### Early leukemic chromatin remodeling at enhancers was consistent between independent clones

Compared to normal progenitors, early leukemic subclones showed a strong chromatin remodeling at enhancers, which was largely maintained during clonal evolution. We identified ∼40,000 enhancer elements based on the density of H3K27ac or H3K4me1 and the absence of strong H3K4me3 enrichment. Many of these enhancers changed the levels of chromatin marks in early leukemic subclones compared to normal progenitors. As an example, two enhancers in the proximity of the *Des* gene showed consistent increases in H3K27ac and H3K4me1 in both clones (Figure 1J). These changes are quantified in a scatterplot (Figure 1K, top) where individual points correspond to the combination of two marks in normal (grey), early leukemic (blue), and more aggressive subclones (red), with proximal gene expression shown as a barplot. *Pitx2* (Figure 1K, middle, and Figure S1K) and *Lmo1* proximal enhancers (Figure 1K, bottom, and Figure S1L) also underwent consistent activation and deactivation of chromatin state. Across the genome, 17,338 enhancers changed H3K27ac or H3K4me1 >2-fold in at least one clone (Figures 1L, S1M). These changes correlated between the two marks and were consistent between two clones (Figure 1L), which shared the majority of enhancers with >2-fold change of H3K27ac and a large fraction of enhancers with >2-fold change of H3K4me1 (Figure 1M). The magnitude of these leukemic changes was largely maintained between two stages of clonal evolution (Figure 1N). The minority of enhancers that showed stage-specific differences are the focus of our further analyses below.

### Clonal evolution was associated with clone-specific changes of gene expression and promoter chromatin states that preferentially reversed early leukemic changes

We then focused on the patterns of chromatin and expression changes upon clonal evolution. As an example, in normal progenitors the *Fhdc* promoter was in a repressed bivalent state with high H3K27me3, modest H3K4me3, and no H3K27ac (Figure 2A). At the early leukemic stage in both clones, it became strongly active, with no H3K27me3 but high H3K4me3 and H3K27ac. However, the further clonal evolution partially reversed this activation by shifting back the levels of chromatin marks and expression at the more aggressive stage (Figure 2A). As an opposite example, the *Fyb* promoter was active in normal progenitors, with high H3K27ac and H3K4me3 and no H3K27me3, and shifted to a repressed bivalent state at the early leukemic stage in both clones (Figure 2B). Upon clonal evolution this strong repression was partially reversed by decreasing H3K27me3 and increasing H3K4me3 and H3K27ac at the more aggressive stage (Figure 2B). These changes are quantified in the scatterplots of H3K27me3 vs H3K4me3 (Figure 2C), with additional examples in Figures S2A-F. Collectively, these changes suggest that evolution towards increased cell growth largely involved modulation of promoter activity on a continuous scale rather than on/off switching between extreme chromatin states.

**Figure 2.**
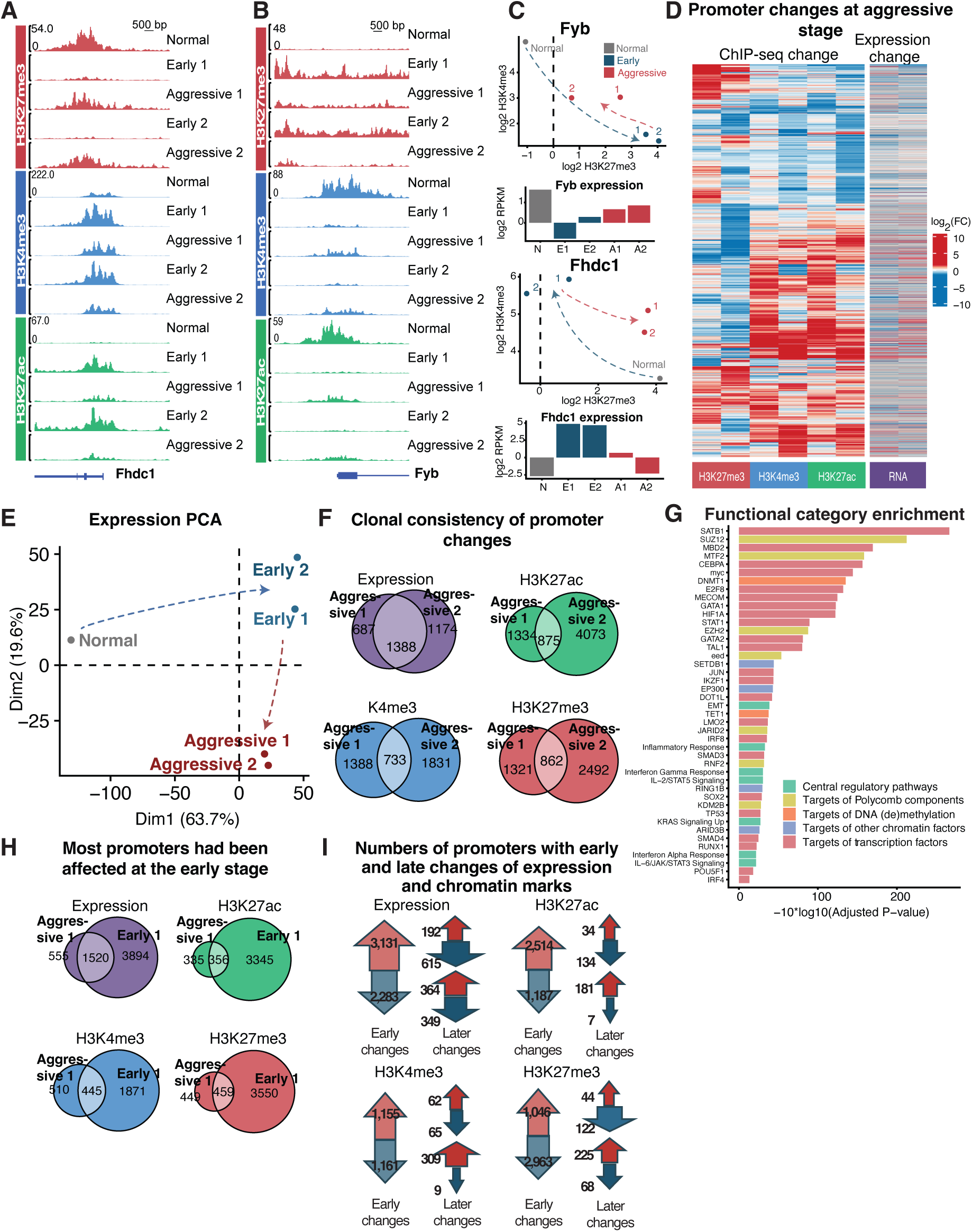
Clonal evolution led to gradual changes of promoter chromatin state and expression which often reversed the early leukemic changes. **A,B.** Examples of ChIP-seq genomic tracks showing promoter chromatin mark and expression changes during clonal evolution. H3K27me3 (red), H3K4me3 (blue), and H3K27ac (green) densities in normal progenitors (top) compared to two independent leukemic clones 1 and 2 at early and aggressive stages. Compared to normal progenitors, Fhdc promoter was activated (**A**) and Fyb promoter was repressed (**B**) to a different degree between the early and aggressive stage of the same clone. **C.** Promoter chromatin state progression for Fhdc and Fyb promoters (tracks shown in **A, B**) in the space of H3K27me3 (x axis) vs H3K4me3 density (y axis). Points represent normal progenitors (grey) and early (blue) and aggressive stages (red) of two leukemic clones (1 and 2). Strong opposite shifts of H3K27me3 and H3K4me3 at the early leukemic stage of both clones (blue arrow) were later partially reversed during clonal evolution (red arrow), concomitant with changes of gene expression (barplots of RNA-seq RPKM). **D.** Heatmap of changes (log2 fold change) of promoter chromatin marks during clonal evolution across all promoters with a substantive absolute level and > 2-fold change of at least one mark, juxtaposed with the changes of gene expression in vivo (log2 fold change, right). Changes were only partially consistent between two independent clones (two columns for each mark and expression). **E.** Principal component analysis (PCA) plot of gene expression in normal progenitors (grey) compared to early (blue) and aggressive stage (red) of two leukemic clones (1 and 2). **F.** Venn diagrams of moderate overlap between two independent clones (Aggressive 1, Aggressive 2) among promoters with changes of gene expression in vivo (purple) and chromatin marks (green, blue, and red) during clonal evolution. **G.** Functional category enrichment among the genes that were consistently differentially expressed upon clonal evolution in both clones in vivo. Bars colored by category types represent adjusted P-value of enrichment according to EnrichR. **H.** Most promoters affected by clonal evolution had undergone prior changes at the early leukemic stage. Venn diagrams of promoters with changes of expression and individual chromatin marks at the aggressive and early stages of the same clone. **I.** Clonal evolution preferentially reversed the prior early leukemic changes at the promoters. Large arrows: numbers of promoters with increase (red) and decrease (blue) of expression and chromatin marks (H3K27ac, H3K4me3, H3K27me3) between normal progenitors and the early leukemic subclone. Smaller arrows near each large arrow: numbers of these promoters that underwent the later increase (red) or decrease (blue) of expression or chromatin mark during clonal evolution. These later changes were skewed in the direction opposite to the early leukemic changes. Data for clone 1; see Figure S2M for clone 2.

Genome-wide, gene expression and chromatin changes during clonal evolution (Figures 2D, S2K) affected many promoters but fewer than at the early leukemic stage (Figure 1G). This smaller yet substantial scale of changes was also evident from principal component analysis (PCA) of gene expression (Figure 2E). The difference between normal progenitors (gray points) and all leukemic subclones regardless of aggressiveness (blue and red) corresponded to the strongest principal component 1 (x axis) that accounted for most of the overall variation (64%). The difference between two stages of clonal evolution (blue vs red) corresponded to a weaker principal component 2 (y axis) and accounted for a smaller proportion of variation (20%).

Compared to the high consistency between independent clones at the early leukemic stage (Figure 1G), changes during the further clonal evolution (Figure 2D) were more clone-specific, suggesting distinct clonal paths. Independent clones 1 and 2 shared 1388 DEGs associated with clonal evolution, which accounted for 54% and 67% DEGs in clones 1 and 2, respectively (Figure 2F, Supplementary Data 1). This overlap was much smaller than ∼85% of shared DEGs at the early stage (Figure 1H). This clonal specificity was even more pronounced among chromatin marks (Figure 2F).

Genes that underwent consistent expression changes in both clones (Figure 2G) were enriched in central regulatory pathways (EMT, interferon and inflammatory responses, IL6/JAK/STAT3, IL2/STA5, KRAS signaling), targets of transcription factors (Satb1, Cebpa, Myc, E2f8, Mecom, Gata1/2, Stat1, Jun, Irf4/8, Smad3/4, p53, Sox2, Oct4/Pou5f1, etc), chromatin modifying proteins (Polycomb components, Dot1l, p300, Arid3b, Setdb1), and DNA methylation and demethylation machinery (DNMT1, Tet1). These factors included general cell cycle regulators (e.g. E2f8), pluripotency factors (Sox2, Oct4, etc), and regulators of leukemic growth (Cebpa, Myc, Dot1l, etc). Among chromatin modifiers, there was a notable enrichment across Polycomb group proteins (Rnf2, Kdm2b, Suz12, Eed, Ezh2, Jarid2, MTF2), suggesting a major role of Polycomb in transcriptional regulation of clonal evolution.

Surprisingly, the majority of DEGs associated with clonal evolution (∼70-75% depending on the clone) had also been differentially expressed earlier between normal progenitors and the early leukemic subclones (Figures 2H, S2L). Consistent with this observation, ∼50% of promoters affected by chromatin remodeling during clonal evolution had also been affected at the earlier leukemic stage (Figures 2H, S2L). However, contrary to our naïve expectation that clonal evolution would further amplify these early leukemic changes, they were preferentially reversed in the aggressive subclones. Figures 2I and S2M schematically show the numbers of promoters with increased and decreased expression at the early stage (large arrows), juxtaposed with the numbers of these promoters that underwent later expression changes at the aggressive stage (small up and down arrows near each large arrow). These changes show the pattern of preferential reversal. For example, among promoters with increased expression at the early leukemic stage (large red arrow), only a fraction was affected upon further clonal evolution (small arrows on the right), but this fraction was preferentially skewed towards downregulation (small blue vs small red arrow). For promoter chromatin marks (Figures 2I and S2M), this preferential reversal of early leukemic changes was even stronger.

Consistent with a strong influence of the bone marrow environment, the differences of gene expression between early and more aggressive subclones in vivo were affected by culturing ex vivo. However, many genes associated with clonal evolution (20% of up- and 13% of downregulated genes) maintained their differential expression regardless of the environment (Figure S2N). We postulated that some of these genes contributed to the cellular memory of the subclonal status, which was maintained ex vivo despite the diminished growth difference between early and aggressive subclones (Figure 1C) and manifested upon secondary re-transplantation (Figure S1D).

Gene expression changes between ex vivo and in vivo in each individual subclone had >50% overlap between the early and more aggressive subclones of the same clone (Figure S2O), consistent with overlapping functional gene categories enriched among these DEGs, which included MYC targets, cholesterol homeostasis, and p53 pathways enriched among genes downregulated in vivo (Figure S2P).

Promoter chromatin changes were correlated with gene expression; the combination of promoter H3K4me3, H3K4me1, H3K27ac, and H3K27me3 differences ex vivo was predictive of expression differences (Figure S2Q, R=0.5). Active mark H3K27ac was the most predictive, followed by H3K27me3 and H3K4me3 (Figure S2Q, heatmap on the right). As expected, chromatin mark changes ex vivo were less predictive of the expression changes in vivo, but active marks H3K27ac and H3K4me3 were still among the most predictive (Figure S2R).

### Chromatin state and expression of direct MLL-AF9 targets were stable throughout clonal evolution

To address whether increased cell growth could be ascribed to increased activity of direct MLL-AF9 targets, we analyzed target promoters with the strongest MLL-AF9 peaks (Homer score ≥ 13) based on published ChIP-seq data in mouse MLL-AF9 HSPCs [27]. Most MLL-AF9 targets were already active in normal progenitors. At the early leukemic stage, 35-40% of these targets (depending on the clone) were further upregulated, while only 5-6% were downregulated, summarized as heatmap (Figure S2G) and trends of combined H3K27me3 and H3K4me3 changes (Figure S2H,I). However, there was no preferential activity shift during clonal evolution, where only a small minority of MLL-AF9 targets were differentially expressed (12-15% depending on the clone) and were mostly downregulated (Figure S2J). Only 3.5-12% of MLL-AF9 targets underwent H3K4me3, H3K27ac, or H3K27me3 changes upon clonal evolution (Figure S2J).

Remarkably, virtually all MLL-AF9 targets that were initially repressed by Polycomb and had high H3K27me3 in normal progenitors underwent a dramatic H3K27me3 depletion and H3K4me3 increase at the early leukemic stage (diagonal arrow towards upper left in Figure S2H,I). These targets included major leukemogenic drivers Hoxa9, Meis1, Six1, Six4, etc (Figures 1E, S1A-E). However, neither their expression nor chromatin state changed further during clonal evolution (Figures 1E,F, S1A-C), with the possible exception of Eya1 and Jmjd1c that changed expression in one of the clones (Figures S1D,E). This absence of clone-consistent changes suggested that direct regulation by MLL-AF9 or major leukemogenic drivers was unlikely to be a primary driver of clonal evolution.

### Clonal evolution preferentially activated chromatin state at enhancers and reversed their early leukemic changes

Clonal evolution was associated with widespread modulation of enhancer chromatin marks. As an example, the Net1 enhancer increased both H3K27ac and H3K4me1 at the more aggressive stage (Figure 3A), whereas the enhancer upstream of gene Zfp516 decreased both marks (Figure 3B, see additional examples in Figure S3A-C). These changes are quantified as scatterplots of H3K4me1 vs H3K27ac at the same enhancer in Figures 3C, S3D-F. Similar to promoters (Figures 2A-C, S2A-F), chromatin mark levels at these enhancers were gradually modulated, as opposed to switching between extreme chromatin states. Globally, clonal evolution was associated with a strong skew towards activation of enhancer chromatin states (Figures 3D, S3G). The consistency of H3K27ac and H3K4me1 changes in two independent clones was significant (20-22% for H3K27ac and 11-15% for H3K4me1 depending on the clone, Figure 3E) but lower than the clonal consistency of chromatin mark changes at promoters (Figure 2F).

**Figure 3:**
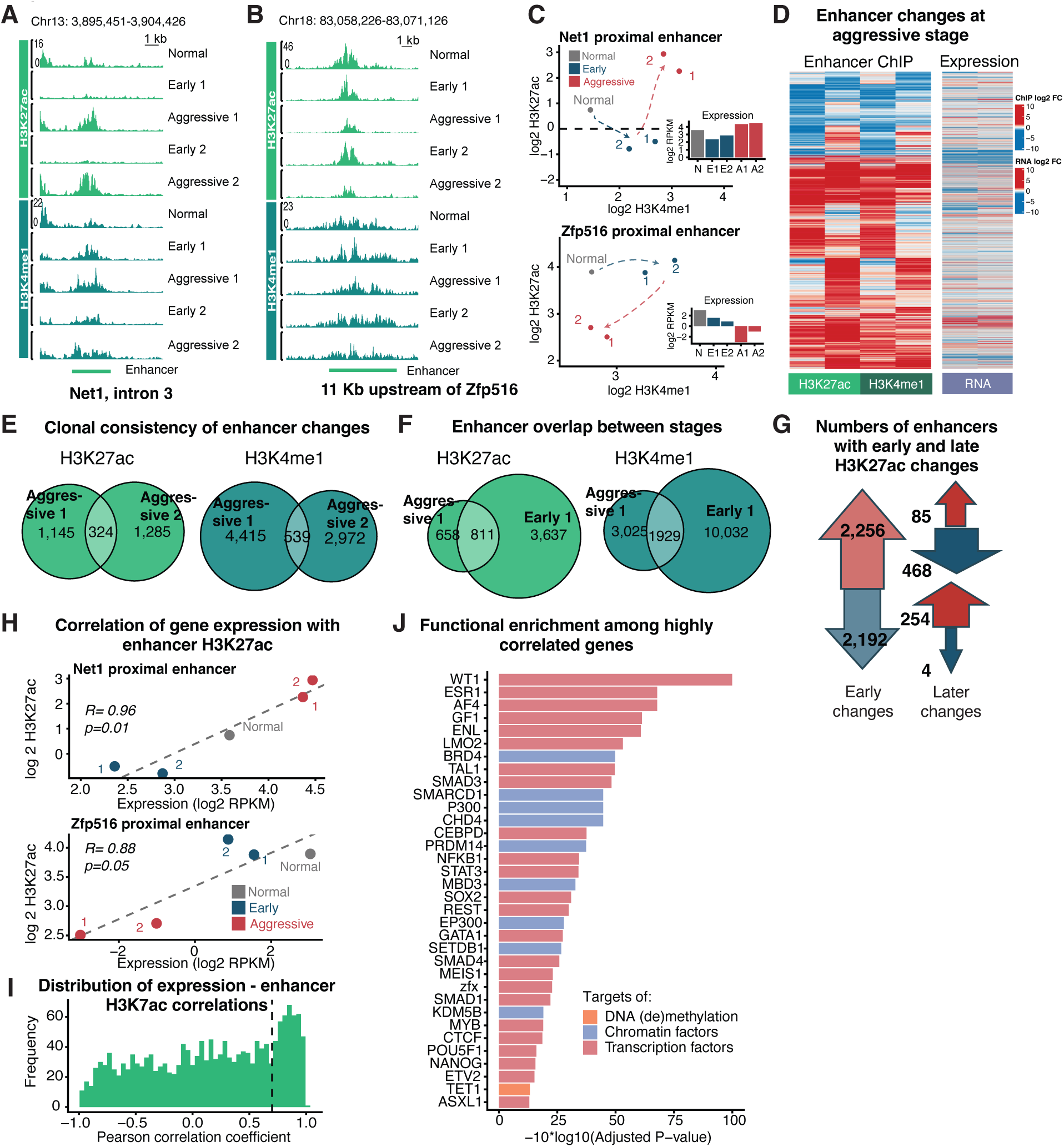
Clonal evolution during preferentially activated enhancer chromatin states and often reversed early leukemic changes at enhancers. **A,B.** Examples of ChIP-seq genomic tracks showing enhancer chromatin mark changes during clonal evolution. H3K27ac (green) and H3K4me1 (dark green) densities in normal progenitors (top) compared to two independent leukemic clones 1 and 2 at the early and aggressive stages. Chromatin state of an intronic enhancer in Net1 gene became more active (**A**) and chromatin state of an enhancer proximal to gene Zfp516 became less active upon clonal evolution in both clones (**B**). **C.** Chromatin state progression in the space of H3K27ac vs H3K4me1 density at enhancers shown in **A, B.** Points represent normal progenitors (grey) and the early (blue) and aggressive stages (red) of two leukemic clones (1 and 2). Changes upon clonal evolution (red arrows) were accompanied by proximal gene expression changes (barplots of RNA-seq RPKM). **D.** Heatmap of H3K27ac and H3K4me1 density changes (log2 fold change) upon clonal evolution across all enhancers with a substantive absolute level and > 2-fold change of at least one mark, juxtaposed with the changes of gene expression in vivo (log2 fold change, right). Changes were only partially consistent between two independent clones (two columns for each mark). **E.** Venn diagrams of moderate overlap between two independent clones (Aggressive 1, Aggressive 2) among enhancers with changes of H3K27ac (green) and H3K4me1 (dark green) upon clonal evolution. **F.** Most enhancers affected by clonal evolution had undergone prior changes at the early leukemic stage. Venn diagrams of enhancers with changes of H3K27ac and H3K4me1 at the aggressive and early stages of the same clone. **G.** Clonal evolution preferentially reversed the prior early leukemic changes of H3K27ac at enhancers. Large arrows: numbers of enhancers with increase (red) and decrease (blue) of H3K27ac between normal progenitors and the early leukemic subclone. Smaller arrows near each large arrow: numbers of these enhancers that underwent the later increase (red) or decrease (blue) of H3K27ac during clonal evolution. These later changes were skewed in the direction opposite to the early leukemic changes. Data for clone 1; see Figure S3I for clone 2. **H.** Enhancer H3K27ac level is often quantitatively correlated with the expression of the proximal gene. Correlation scatterplots of enhancer H3K27ac density vs proximal gene expression across normal progenitors (grey), early (blue) and aggressive leukemic stages (red) for two enhancers shown in **A-C**. Pearson correlation coefficient and linear regression P-value are indicated. **I.** Enhancers increasing H3K27ac during clonal evolution show preferential positive correlation of H3K27ac level with proximal gene expression. Histogram of all Pearson correlation coefficients between enhancer H3K27ac and gene expression (calculated as in **H**) is strongly skewed towards higher R and includes a distinct enhancer subset with strong correlation (R > 0.7, dotted line). **J.** Functional category enrichment among genes proximal to the enhancer subset shown in **I**. Bars colored by category types represent adjusted P-value of enrichment according to EnrichR.

Like promoters (Figure 2H), a large proportion of enhancers affected by clonal evolution had already undergone changes at the early leukemic stage. This proportion was especially large for H3K27ac (Figures 3F, S3H). Surprisingly, most of these enhancers reversed their initial H3K27ac changes. Figures 3G and S3I schematically show this reversal by the numbers of enhancers with increased and decreased H3K27ac at the early stage (large arrows), juxtaposed with the numbers of the same enhancers that underwent later H3K27ac changes at the aggressive stage (small up and down arrows near each large arrow). For example, among enhancers with increased H3K27ac at the early leukemic stage (large red arrow), only a fraction was affected upon further clonal evolution (small arrows on the right), but this fraction was preferentially skewed towards H3K27ac decrease (small blue vs small red arrow).

These changes of enhancer chromatin marks were associated with various transcription factor binding motifs according to MEME[28] (Figure S3J). Transcription factors associated with the coordinated changes of both H3K27ac and H3K4me1 included ETS family proteins and related PU.1 factor that have been implicated in hematopoiesis and leukemogenesis [29,30]; CEBPB/D/E/G involved in myeloid hematopoiesis and aggressive leukemia [31–33]; and ZNF384 with reported leukemic fusions and regulation of hematopoietic stem cell factors[34]. Motifs associated with only H3K27ac changes involved major regulatory factors STAT2, BCL11B, IKZF1, NR5A1/2, and KLF9, whereas H3K4me1 changes were associated with leukemic regulators CEBPA, MYB, RUNX1, pluripotency factor KLF4, AP-1 factor JUN, and other factors (Figure S3J). Taken together, enhancer chromatin remodeling during clonal evolution was associated with motifs for ETS, CEBP, AP-1 factors and other differentiation and leukemic regulators.

For each enhancer, we calculated correlation between its H3K27ac level and the expression of the proximal gene across the five profiled cell types (normal progenitors and two stages of evolution in two independent leukemic clones). As an example, Figure 3H quantifies these correlations for two enhancers depicted in Figures 3A,B. Genome-wide, enhancers that underwent H3K27ac changes during clonal evolution had a remarkable skew towards high correlation with proximal gene expression (histogram in Figure 3I), suggestive of transcriptional regulation. We focused on 406 enhancers with the highest correlation to gene expression (Pearson R>0.7, right of the dotted line in Figure 3I). EnrichR analysis of the corresponding 307 genes (Figure 3J) revealed strong enrichment in the targets of major differentiation, hematopoiesis, and leukemogenesis regulators (Wt1, Esr1, Lmo2, Gf1, Myb, Tal1, Cebpd, Stat3, Meis1), components of super-elongation complex also known as MLL fusion partners (Af4/Mllt3 and Af9/Enl), pluripotency factors (Sox2, Nanog, Pou5f1, Prdm14), and chromatin modifiers (Brd4, p300, Smarcd1, Setdb1) often implicated in hematopoiesis and leukemia. The enrichment of Nfkb1 and Smad3/4 targets suggested the involvement of NFKB and TGFβ pathways. Although many enriched categories were similar to the ones among the whole population of genes affected by clonal evolution (Figure 2G), there were notable differences. Polycomb components were not as strongly enriched (Figure 3J), suggesting lesser involvement of Polycomb control; whereas targets of Kdm5b and Smad1 were enriched exclusively among enhancer-correlated promoters, suggesting that Kdm5b, an H3K4 histone demethylase known to suppress AML aggressiveness [15] and Smad1, a BMP pathway effector, are involved in enhancer-mediated gene regulation.

Taken together, these changes of gene expression and chromatin marks at promoters (Figures 2A,B,H,I) and enhancers (Figures 3A,B,F,G) during clonal evolution suggest a few important observations. First, most of these changes were gradual quantitative shifts rather than binary switches between silent and active states. These gradual shifts were often consistent between clones to a considerable degree of quantitative detail. Second, most of these shifts were preceded by the prior changes of the same promoter or enhancer at early leukemic stage. When compared to normal progenitors, these genes most often maintained the early direction of shift, either activation or repression, but altered the magnitude of this change (Figure 1I,N). This suggests that clonal evolution involved quantitative adjustment of initial dysregulation that had occurred during leukemogenesis. Unexpectedly, this quantitative adjustment often dampened this initial dysregulation. As one possible explanation of fitness advantages conferred by this dampening, some of the extremely strong downstream effects of initial leukemogenic lesion (MLL rearrangement) may require fine tuning or even toning down for further increase of cell growth and survival. This fine tuning might be mediated via additional mutagenesis, changes in expression of key transcription factors, chromatin remodeling by epigenetic factors, or other mechanisms.

### Core genes defined by clonally consistent expression and chromatin changes in vivo and ex vivo

We next aimed to leverage the observed variability between clones and conditions to filter out the genes that show only clone-specific or condition-specific changes and focus on the genes that were consistently affected by clonal evolution in both clones (Figure 2F) and maintained their differential status both in vivo and ex vivo (Figure S2N). We hypothesized that this consistent gene set would be enriched in the genes that causally regulate cell growth during clonal evolution in vivo and maintain the memory of the subclone’s evolutionary stage through culturing ex vivo. Specifically, we defined a core set of genes with consistent expression and promoter chromatin changes persisting between ex vivo and in vivo conditions in both independent clones (Figure 4A). As a result, we identified 17 such genes that were consistently upregulated and 11 genes that were consistently downregulated during clonal evolution (Figures 4B, S4A).

**Figure 4.**
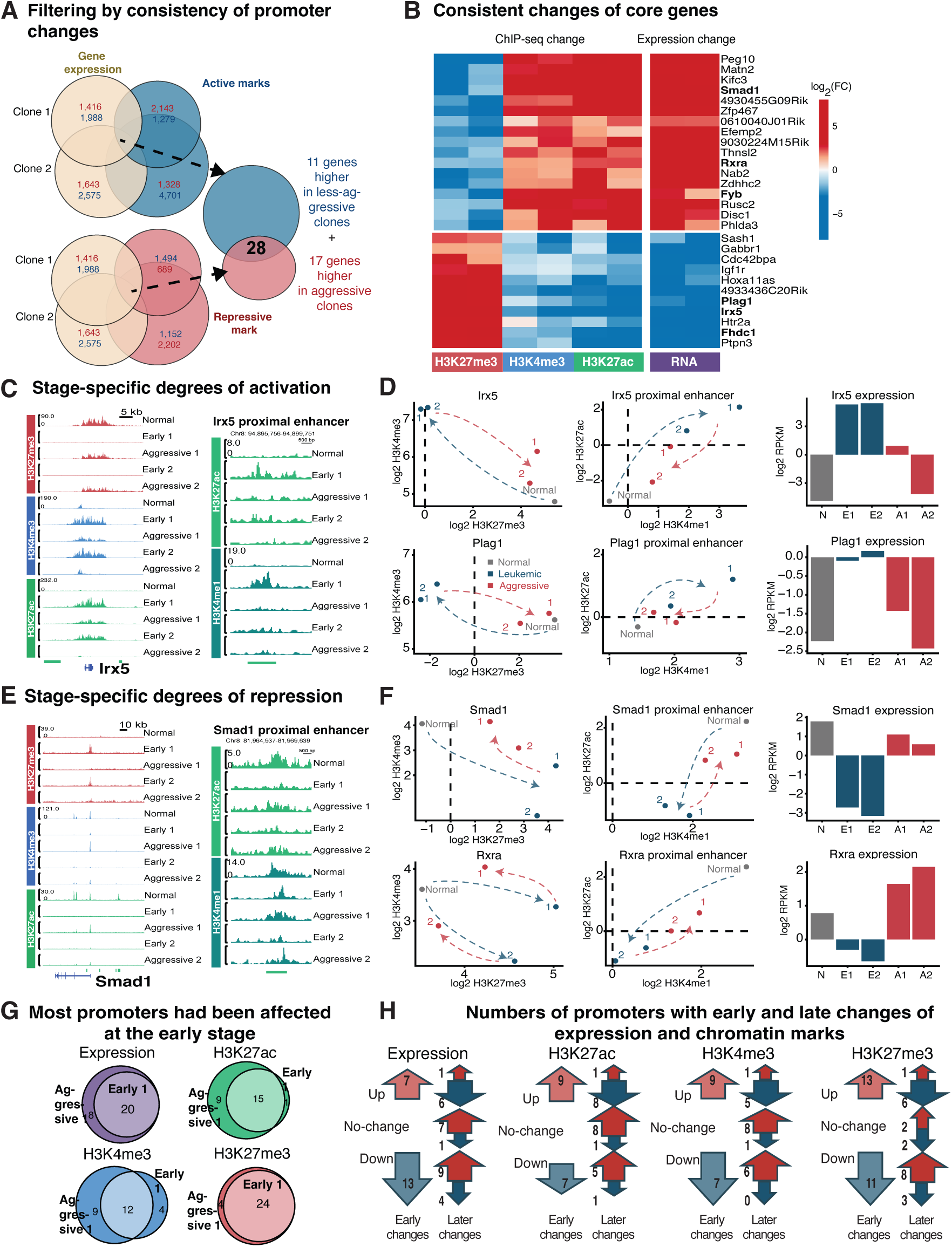
Core genes with consistent changes of promoter chromatin state and expression in both independent clones. Filtering genes by the consistency of changes of gene expression in vivo (yellow) together with active H3K27ac and H3K4me3 (blue) and repressive H3K27me3 (red) promoter marks ex vivo during clonal evolution in both clones. Numbers of promoters with expression or chromatin mark increase and decrease are indicated in red and blue, respectively. A core set of 28 candidate genes had fully consistent behavior of expression and chromatin marks in both clones. **B.** Heatmap of promoter H3K27me3, H3K4me3, H3K27ac and expression changes (fold change in log2 scale) for the core gene set during clonal evolution of two clones. **C.** Examples of genomic ChIP-seq tracks at Irx5 (left) and its proximal enhancer (right) where the magnitude of the early leukemic activation was later adjusted during clonal evolution. **D.** Examples of promoter chromatin state progression in the space of H3K27me3 (x axis) vs H3K4me3 (y axis) for core genes Irx5 (tracks in **C**) and Plag1 (tracks in Figure S5B). Points represent normal progenitors (grey) and early (blue) and aggressive stages (red) of two leukemic clones (1 and 2). H3K27me3 loss and H3K4me3 gain at the early leukemic stage of both clones (left plots, blue arrows) were later partially reversed upon clonal evolution (left plots, red arrows), consistent with in vivo expression changes (right barplots). These changes were paralleled by H3K27ac and H3K4me1 at a proximal enhancer (middle plots). **E.** Examples of genomic ChIP-seq tracks at Smad1 (left) and its proximal enhancer (right) where the magnitude of the early leukemic repression was later adjusted during clonal evolution. **F.** Examples of promoter chromatin state progression in the space of H3K27me3 vs H3K4me3 for core genes Smad1 (tracks shown in **E**) and Rxra (tracks in Figure S5C). H3K27me3 gain and H3K4me3 loss at the early leukemic stage of both clones (left plots, blue arrows) were later partially reversed upon clonal evolution (left plots, red arrows), consistent with in vivo expression changes (right barplots). These changes were paralleled by H3K27ac and H3K4me1 at a proximal enhancer (middle plots). **G.** Virtually all core genes had undergone prior changes at the early leukemic stage. Venn diagrams of promoters affected by the changes of expression and chromatin marks at aggressive and early stages of the same clone (second clone shown in Figure S4D). **H.** Clonal evolution preferentially reversed the prior early leukemic changes at the promoters of core genes. Large arrows: numbers of promoters with increase (red) and decrease (blue) of expression and chromatin marks (H3K27ac, H3K4me3, H3K27me3) between normal progenitors and the early leukemic subclone. Smaller arrows near each large arrow: numbers of these promoters that underwent the later increase (red) or decrease (blue) of expression or chromatin mark during clonal evolution. These later changes were skewed in the direction opposite to the early leukemic changes. Data for clone 1; see Figure S4E for clone 2.

While none of these genes were strong MLLr targets, many were known regulators of hematopoiesis, leukemia and other cancers. Among upregulated genes, Rxra is a hormone receptor mutated and therapeutically targeted in an aggressive form of AML, acute promyelocytic leukemia [35]. Smad1 is a key effector in BMP signaling pathway playing an important role in normal hematopoiesis [36,37] and leukemia [38,39]. Peg10 is known to enhance tumor growth by driving cell cycle progression through the inhibition of TGFβ pathway [40,41]. Consistent with this mechanism, another upregulated gene Matn2 is activated by TGFβ signaling inhibition [42].

Among downregulated genes, homeobox factor Irx5 plays major roles in early development [43,44], metabolism [45] and cancers [46,47], and is upregulated in AML patients and cell lines [48], but no mechanistic role in AML has been reported. Irx5 is a strong target of MLL-AF9 binding and transcriptional regulation [49] and was dramatically activated at the early leukemic stage (Figures S4A, 4C,D). However, in contrast to most MLL-AF9 targets (Figures S2G-I), Irx5 later reversed this activation and was downregulated upon clonal evolution (Figure S4A, 4B-D). Plag1 is an oncogene involved in the origin of leukemia [14,50] and was upregulated at the early leukemic stage (Figures S4A,B, 4D). However, similar to Irx5, Plag1 later reversed this activation and was downregulated upon clonal evolution (Figures S5A,B, 4B,D). To our knowledge, Plag1 has not been previously implicated in regulation of clonal evolution. Protein tyrosine phosphatase Ptpn3 is a potential tumor suppressor facilitating the activity of TGFβ pathway by stabilizing TGFβ receptor [51]. Ptpn3 downregulation was in line with the upregulation of Peg10 and Matn2 associated with TGFβ inhibition (Figure 4B). Finally, Igf1r is a major regulator of cell transformation and oncogenesis, including AML [52], consistent with Igf1r upregulation at the early leukemic stage (Figure S4A). However, once MLLr leukemia is established, Igf1r is no longer required for clonal propagation [52], in line with the observed Igf1r reversal towards downregulation during later clonal evolution (Figures S4A, 4B).

### Gradual modulation of core genes involved reversal of early leukemic changes

Compared to normal progenitors, most core genes maintained their activation or repression at both leukemic stages, but its magnitude was stage-specific (Figure S4A). Of note, enhancers proximal to these genes often had similar gradual shifts of chromatin state. As an example, strong Polycomb repression of *Irx5* promoter in normal progenitors was completely removed at the early leukemic stage but was partially reversed upon clonal evolution by increasing H3K27me3 to a moderate level while partially decreasing H3K4me3 and H3K27ac (Figure 4C, left). This partial reversal of early activation was paralleled by H3K27ac and H3K4me1 changes at a proximal enhancer (Figure 4C, right). This reversal is quantified in Figure 4D as scatterplots of promoter H3K27me3 vs H3K4me3 (left) and enhancer H3K4me1 vs H3K27ac (middle), along with gene expression changes (barplots on the right). *Plag1* had an even stronger reversal of early promoter and enhancer activation, which resulted in an almost complete return to the chromatin state and expression observed in normal progenitors (Figures S4B, 4D). This return suggests that despite the known oncogenic role of Plag1 [14,50], its upregulation may paradoxically be absent in patients with advanced AML.

Conversely, promoters and proximal enhancers of *Smad1* (Figure 4E) and *Rxra* (Figure S4C) were active in normal progenitors, followed by strong repression at the early leukemic stage and its partial reversal during clonal evolution. These changes are quantified in Figure 4F. As RXRA is a well-known leukemogenic driver in a different genetic background of PML-RARA fusion [35], its behavior in MLL-AF9 clones may suggest a more general role in driving cell growth during clonal evolution.

The fraction of the core genes that had already been affected at the early stage (Figures 4G, S4D) was even larger than the genome-wide fraction among all promoters affected by clonal evolution (Figures 2H, S2L). Most of the core promoters reversed their early leukemic changes of gene expression and chromatin marks (Figure 4H, S4E), rather than further amplifying these early changes.

### Functional validation of candidate growth regulators

From these core genes, we selected Irx5, Plag1, and Smad1 for validation as candidate regulators of cell growth in leukemic progenitors. This selection was based on the magnitude and clonal consistency of chromatin mark and expression changes (Figures 4B,C-F). Since transcription factors Irx5 and Plag1 were consistently repressed upon clonal evolution (Figures 4C,D, S4B), we hypothesized that they were suppressors of leukemic cell growth. Since BMP pathway effector Smad1 was consistently activated (Figures 4E,F), we hypothesized that Smad1 is a driver of leukemic cell growth.

To validate phenotypic effects of Irx5, Plag1, and Smad1, we used lentiviral transduction to overexpress these genes in leukemic progenitor clones. Irx5 and Plag1 were overexpressed in more aggressive leukemic progenitors to test their suppressive effect while Smad1 was overexpressed in the earlier, less aggressive leukemic progenitors to test its activating effect. As a positive control, the known oncogenic driver, Kras G12V mutant [53], was overexpressed in the same earlier leukemic progenitors. Expression was confirmed at the RNA level (Figure 5A). Overexpression of these genes had a strong effect on cell growth (Figure 5B). Smad1 overexpression in an early leukemic subclone increased growth rate by 45% (decreased the doubling time from 13.7 to 9.5 hours), even more than Kras G12V expression (decreased the doubling time from 13.7 to 11.7 hours). The overexpression of Irx5 and Plag1 in an aggressive subclone had strong effects in the opposite direction, suppressing growth rate by 62% (increase of doubling time from 10.1 to 26.2 hours) and 48% (increase of doubling time from 10.1 to 19.3 hours), respectively (Figure 5B).

**Figure 5.**
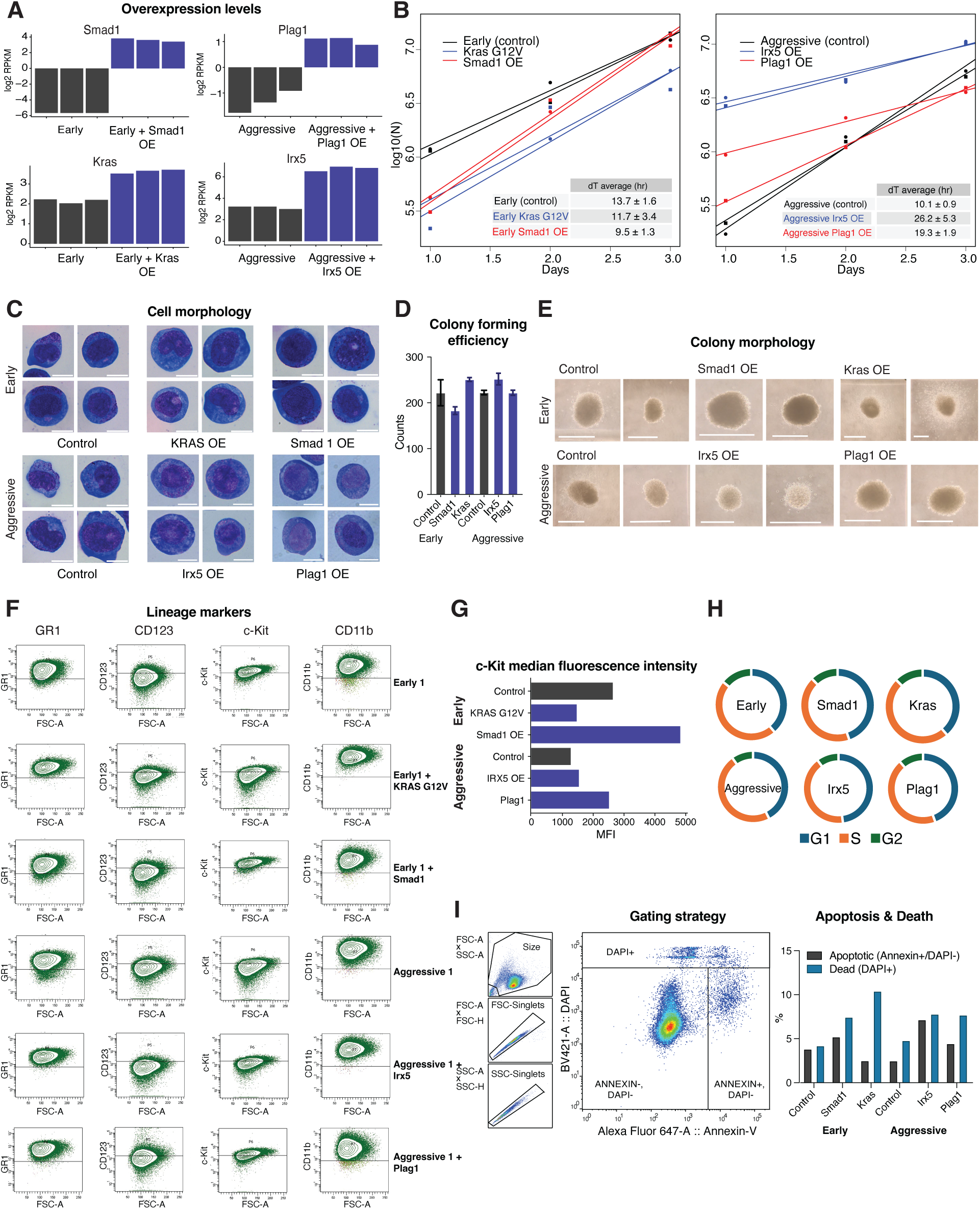
Effects of overexpressing selected core genes validate their causal roles. **A.** Smad1 and Kras G12V mutant as a control were overexpressed in the early stage subclone, whereas Irx5 and Plag1 were overexpressed in the advanced more aggressive subclone of the same leukemic clone. Barplots of RNA-seq gene expression values (RPKM) in transduced compared to control untransduced cells (n=3 biological replicates). **B.** Overexpression of Smad1 increased, whereas overexpression of Irx5 and Plag1 decreased cell growth. Ex vivo cell growth curves and the corresponding doubling times for transduced cells compared to untransduced controls. Data are represented as mean +/- SEM. **C.** Representative images of Wright-Giemsa–stained cells transduced with each gene, compared to control. Scale bar, 1 micron. **D.** Colony forming efficiency by transduced clones did not show strong changes compared to control. Barplot of colony numbers in methylcellulose (mean and SD, n=2) per 1500 cells. **E.** Examples of individual colonies formed by the transduced cells compared to the control early and aggressive subclones. **F.** Flow cytometry analysis of c-Kit, Gr1, CD123, and CD11b expression in each transduced subclone compared to control suggest no strong changes of myeloid hematopoietic lineage. **G.** Median c-Kit fluroscence intensity in the transduced cells compared to the control early and aggressive subclones. **H.** Fractions of cell cycles phases (G1, S, G2/M) for the transduced cells compared to the control early and aggressive subclones. **I.** Annexin V apoptosis assay. Left, gating strategy (example). Right, barplot of fractions of apoptotic (Annexin+ / DAPI-) and dead cells (DAPI+).

The size and morphology of leukemic cells overexpressing either of these genes were similar to non-transduced control cells by Wright-Giemsa staining (Figure 5C). We also assessed effects of gene overexpression on colony formation using a growth in methylcellulose assay (Figures 5D,E). The more aggressive subclone had similar colony formation capacity to the early subclone, and gene overexpression did not have a dramatic effect on colony numbers (Figure 5D). However, Irx5, Plag1, and Smad1 had different effects on the individual colony growth (Figure 5E). Colonies formed by a more aggressive subclone, as opposed to the early subclone, often had a pattern of peripheral spreading. Irx5 or Plag1 overexpression strongly reduced this spreading, and Irx5 overexpression in particular led to visually lower cell numbers per colony, consistent with a strong decrease of cell growth. Kras overexpression in the early subclone strongly increased this pattern of spreading. Smad1 overexpression did not increase the spreading but produced much denser colonies with visually larger numbers of cells per colony, consistent with a strong increase of cell growth.

The early progenitor marker IL-3R (CD123) and markers of mature myeloid cells Gr-1 and CD11b remained stable when compared to non-transduced leukemic progenitors (Figures 5F, S5A-C), suggesting no changes in myeloid lineage. To understand the effects on leukemia initiating cell potential, we quantified the intensity of c-Kit (CD117) staining (Figures 5F,G). Compared to the early clone, the more aggressive subclone had a decreased stem cell potential (22% vs 73%). In the early subclone, Smad1 overexpression further increased stemness from 73% to 97%, whereas Kras overexpression decreased stemness from 73% to 29%. In the aggressive subclone, Irx5 overexpression caused only a modest change of stemness from 22% to 25%, whereas Plag1 overexpression increased stemness from 22% to 70% (Figure 5G).

To analyze the effects on cell cycle, we performed flow cytometry using propidium iodide DNA staining in transduced and non-transduced progenitors (Figures 5H, S5D,E). The overexpression of Smad1, Irx5, and Plag1 resulted in only modest shifts between the phases of cell cycle. In the early subclone, Smad1 overexpression decreased S-phase fraction by 5%, from 48% to 43%, and increased G1 fraction by 5%, from 39% to 44%, suggesting a moderate extension of G1 at the expense of S-phase. In the aggressive subclone, both Irx5 and Plag1 overexpression produced similar small shifts towards the extension of G1-phase and decrease of S-phase (Figures 5H, S5D,E).

To quantify apoptosis, we performed flow cytometry of transduced and non-transduced leukemic progenitors using Annexin V staining in combination with the DAPI cell viability dye (Figure 5I). The fraction of apoptotic (Annexin+ / DAPI-) cells was small, with the lowest fraction of ∼2% apoptotic cells observed in non-transduced aggressive subclone and in Kras-overexpressing early subclone, and the highest fraction of ∼7% apoptotic cells observed in Irx5-overexpressing aggressive subclone. These modest differences were consistent with the effects of Kras and Irx5 on cell growth, although the absolute fractions of apoptotic cells were low and therefore unlikely to fully account for the observed growth effects.

### Convergence of Smad1, Irx5, and Plag1 downstream transcriptional effects suggests a common regulatory network

To understand transcriptional effects of Smad1, Irx5 and Plag1 overexpression, we performed RNA-seq and compared genome-wide differential expression patterns (Figure 6A-C, Supplementary Data 2). Consistent with their opposite effects on cell growth (Figure 5B), Smad1 effects were largely opposite to the effects of Plag1 and Irx5 (Figure 6A). These opposite effects were associated with targets of key regulatory proteins (Figure 6B): Polycomb components Kdm2b, Ring1b, Suz12, Ezh2 and Mtf2, central chromatin factors Brd4, Setdb1 and Arid3a, DNA methylation and demethylation components Dnmt1 and Tet1, and major transcription factors (Myc, Myb, Irf8, Ikzf1, Wt1, Rxr, Tal1, Sox2, Nanog, p53, Smad3 etc). There was a significant overlap between individual DEGs affected in the same direction by Irx5 and Plag1 (Figures 6C); for example, both upregulated Bcorl1, a Polycomb component strongly involved in leukemia [58,59] and downregulated Lin28b, which regulates hematopoiesis by transcriptionally inhibiting Polycomb component Cbx2 [60] (heatmap in Figure 6D and schematic in Figure 6E). Notably, Smad1 expression itself was downregulated by Plag1 overexpression (Figure 6D), suggesting a negative transcriptional regulation of Smad1 by Plag1 (Figure 6E). A significant fraction of Irx5 and Plag1 targets was affected by Smad1 in the opposite direction (Figures 6C). For example, Kdm5b, Glis2, Smad3 and DNAjc6 were upregulated by Irx5 and Plag1 and downregulated by Smad1 (Figure 6E). H3K4me2/3 demethylase Kdm5b is a potent suppressor of AML aggressiveness [15]. Glis2 is a developmental transcription factor implicated in the control of AML cell differentiation [61]. Smad3 is a central mediator of signals from TGFβ-like proteins widely implicated in leukemia [62,63], whereas Smad1 mediates signals from BMP proteins [38]. Downregulation of Smad3 by Smad1 suggests an interesting negative crosstalk between BMP and TGFβ signaling, reminiscent of reported TGFβ mediated repression of BMP targets in various contexts [64]. Together with Smad3 upregulation by Irx5 and Plag1, this implies opposite roles of BMP and TGFβ pathways in controlling leukemic clonal growth. Meis2, oncogenic factor in MN1-induced leukemia and AML1-ETO AML [65], and Bmx, a Tec kinase contributing to drug resistance in AML [66], were downregulated by Irx5 and Plag1 and upregulated by Smad1 (Figure 6D,E).

**Figure 6.**
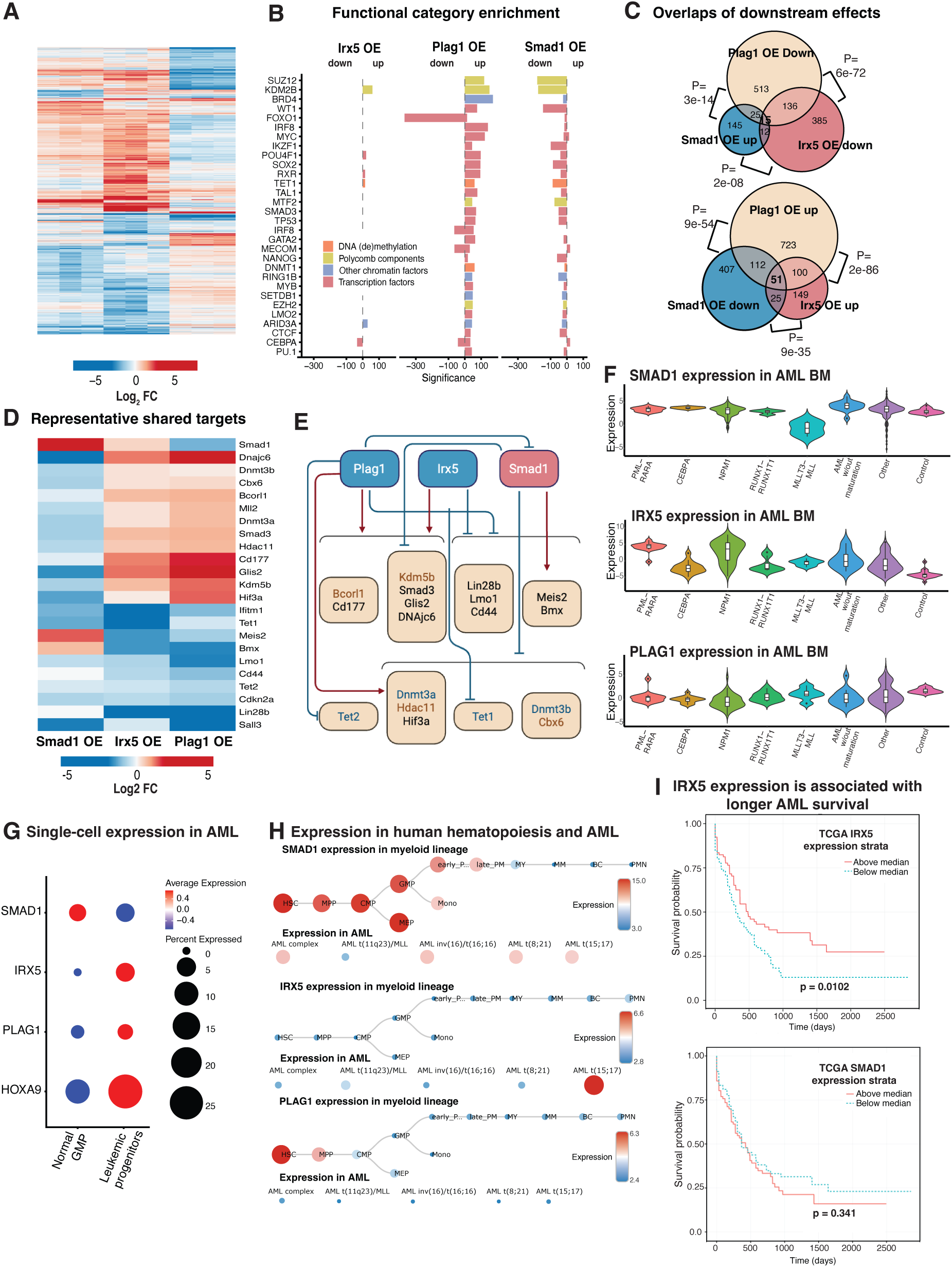
The roles of Smad1, Irx5, and Plag1 are supported by convergent downstream effects on a common regulatory network and by AML patient data. **A.** Genome-wide transcriptional changes caused by Smad1 overexpression in mouse leukemic progenitors are largely opposite to the changes caused by Irx5 and Plag1 overexpression. Heatmap of RNA-seq expression changes compared to control (log2 fold change) among genes differentially expressed (> 2 fold change, FDR < 0.05) in any of the three transduction experiments. **B.** Overlap of functional categories enriched among genes up- and downregulated (bars to the right and left of dotted line, respectively) by Irx5, Plag1, and Smad1 overexpression. Bars colored by category types represent 10log10 of adjusted P-value of enrichment according to EnrichR. **C.** Overlaps between downstream transcriptional effects of the three overexpressed genes. Venn diagrams of the genes downregulated by Plag1 and Irx5 and upregulated by Smad1 (top), and of the genes upregulated by Plag1 and Irx5 and downregulated by Smad1 (top). Hypergeometric P-values for pairwise overlaps are indicated. **D.** Heatmap of RNA-seq expression changes compared to control (log2 fold change) for selected downstream targets consistently affected by Smad1, Irx5, and Plag1 overexpression. **E.** Smad1, Irx5, and Plag1 effects suggest a common downstream regulatory network (schematic). **F.** Violin plots showing the distributions of SMAD1, IRX5, and PLAG1 expression (log2 scale) within various AML subtypes compared to healthy control group (pink, right). SMAD1 is downregulated in MLL-rearranged AMLs. IRX5 is upregulated across major AML subtypes. PLAG1 expression in MLL-rearranged AMLs is not significantly different from normal control and modestly downregulated in some other AML subtypes. **G.** Analysis of previously published single-cell RNA-seq data in granulocyte/macrophage progenitors (GMPs) from AML patients [16] confirms lower expression of SMAD1 and higher expression of IRX5 and PLAG1 in human leukemic progenitors. These differences are smaller but similar in scale to the expression differences for a major leukemic driver HOXA9. **H.** SMAD1, IRX5, and PLAG1 expression levels in public data from normal human myeloid lineage (GEO GSE42519) and various AML types (GEO GSE13159), analyzed in BloodSpot. SMAD1 is highly expressed whereas IRX5 and PLAG1 are silent in normal human GMPs. SMAD1 is downregulated whereas IRX5 is upregulated specifically in MLLr AML t(15;17). PLAG1 expression is low across AML types. **I.** Analysis of TCGA AML patient cohort implemented in BloodSpot. Kaplan-Meier curves for AML patients stratified by IRX5 and SMAD1 expression. High IRX5 expression is significantly associated with longer overall survival in AML patients (top), while higher Smad1 expression tends to correspond to poorer survival, albeit with a non-significant P-value (bottom).

Smad1 overexpression downregulated central DNA methylation and chromatin remodeling factors, which were often co-regulated by Irx5 or Plag1 (Figure 6E, lower row). For example, DNA methylase Dnmt3a and histone deacetylase Hdac11 were downregulated by Smad1 and upregulated by Plag1; whereas key DNA demethylation components Tet1 and Tet2 were downregulated by Irx5 and Plag1, respectively. DNA methylation and demethylation machineries, especially DNMT3A and TET2 are strongly involved in myeloid malignancies including AML [7,67,68], consistent with the enrichment of DNA methylation and demethylation targets among genes affected by clonal evolution (Figure 2G) and by Irx5, Plag1 and Smad1 overexpression (Figure 6B).

Collectively, downstream effects of Smad1, Irx5, and Plag1 converged on key transcription factors (Figure 6E, black), chromatin remodelers (Figure 6E, red), and components of DNA methylation machinery (Figure 6E, blue). Clonal growth driver Smad1 tended to downregulate repressive epigenetic factors (Polycomb proteins, histone deacetylases and demethylases, de novo DNA methylases), whereas clonal growth suppressors Irx5 and Plag1 tended to have opposite effects. Involvement of these epigenetic repressors in the origin and maintenance of leukemia has been emerging in the literature [67], and our results suggest their involvement in leukemic clonal evolution. At least one of these modifiers, histone H3K4 demethylase Kdm5b, is a direct suppressor of AML aggressiveness[15], suggesting that the broader set of Irx5, Plag1, and Smad1 targets may include new candidate regulators of leukemic clonal evolution.

### AML patient data support clinical relevance of identified gene regulators

As our predictions of leukemic cell growth regulators were based on the analysis of clonal evolution in a relatively artificial model of mouse MLL-AF9 overexpressing cells, we asked whether our predictions are applicable to AML in humans. For that we analyzed public gene expression and survival data in AML patient cohorts. These patient data were consistent with gene expression patterns observed in mouse cells and confirmed the role of IRX5 and SMAD1 as markers of AML status in patients. This suggested that our multiomic analysis of clonal evolution in a tractable but artificial mouse model is a valuable predictive method transferrable to human AML.

To assess the expression of SMAD1, IRX5, and PLAG1 across various AML subtypes (NPM1, PML-RARA, CEBPA, etc.) we analyzed the BeatAML 2 database [57] expression data in bone marrow of AML patients, grouped by driver mutations and compared to healthy bone marrow controls (Figure 6F). Assuming that the majority of expression signal in AML bone marrow comes from aggressive leukemic clones, we analyzed expression differences between AML patients and healthy controls and compared them to the differences that we observed in mouse between aggressive leukemic subclones and normal progenitors shown Figures 4D,F.

SMAD1 expression in the BeatAML database is substantially downregulated in MLLr AML patients compared to healthy controls (Figure 6F). This is consistent with our findings in mouse cells, where SMAD1 was strongly downregulated in early leukemic subclones, and despite a partial reversal of this downregulation upon clonal evolution, SMAD1 expression still remained lower than normal (Figure 4F). In other AML subtypes, however, SMAD1 expression is affected to a smaller extent, possibly suggesting a special role of SMAD1 regulation in MLLr AMLs. In contrast, IRX5 is upregulated in MLLr AMLs (Figure 6F). This is consistent with our findings that IRX5 is strongly upregulated in early mouse leukemic subclones, and although this upregulation is partially reversed upon clonal evolution, the level of IRX5 expression remains higher than normal (Figure 4D). The upregulation of IRX5 is also evident across other major AML subtypes, suggesting a more universal IRX5 regulation in various AMLs (Figure 6F). PLAG1 shows no significant expression change among MLLr AML patients and a modest downregulation in some other AML subtypes (Figure 6F). Our observations in mouse progenitors suggest that PLAG1 is strongly upregulated in early mouse leukemic clones, consistent with previously published evidence of its oncogenic role in AML [14,50]. However, during later clonal evolution PLAG1 undergoes near-complete reversal and returns to a very low expression similar to normal progenitors (Figure 4D). Similar reversal during human clonal evolution might contribute to the lower average expression in the bone marrow of AML patients (Figure 6F).

Our observations in mouse leukemic progenitors were also consistent with SMAD1, IRX5, and PLAG1 expression in human leukemic progenitors from MLLr AML patients profiled with single-cell RNA-seq [16]. SMAD1 was downregulated while IRX5 and, to a smaller extent, PLAG1 were upregulated in leukemic compared to normal GMPs (Figure 6G). The magnitude of these differences was somewhat lower but on a similar scale to the dysregulation of a major leukemic driver HOXA9.

According to the BloodSpot database [56], in normal human hematopoiesis SMAD1 is strongly expressed in HSCs and hematopoietic progenitors but is dramatically downregulated during further myeloid differentiation (Figure 6H). SMAD1 is also downregulated in various AML subtypes, especially MLLr AML (Figure 6H), consistent with Smad1 repression in mouse (Figures 4C,E) and human leukemic progenitors, Figure 6G). IRX5 has low expression levels across human HSCs, progenitors, and differentiated cells and in most AML subtypes except for t(15;17) PML-RARA fusion leukemia where IRX5 is strongly upregulated. PLAG1 is strongly expressed in human HSCs and gradually downregulated through stages of myeloid differentiation. Its expression is relatively low in all AML subtypes profiled in BloodSpot (Figure 6H).

Furthemore, survival analysis of public TCGA (https://www.cancer.gov/tcga) AML dataset implemented in BloodSpot[56] revealed the patterns consistent with patient gene expression data (Figure 6H). Higher IRX5 expression had a significant association with longer overall survival (Figure 6A), consistent with Irx5 downregulation during clonal evolution in mouse progenitors (Figure 4C,D) and cell growth suppression upon Irx5 overexpression (Figure 5B). Consistent with Smad1 upregulation during clonal evolution in mouse progenitors (Figure 4E,F) and Smad1 effect on cell growth (Figure 5B), higher SMAD1 expression tended to correspond to shorter overall survival (Figure S5F), albeit the difference was not statistically significant. PLAG1 expression did not show significant survival association with patient survival (Figure 6I).

## DISCUSSION

Clonal evolution in blood cancers is driven by the selection of advantageous genetic and epigenetic alterations [1,5]. Genomic analyses suggest that although the origination of myeloid malignancies involves a relatively small number of mutations [69], the clonal evolution is heterogeneous and can be driven by various additional genetic lesions in individual subclones [1,2,5] during unperturbed clonal evolution or as adaptation to therapy [3,4]. Disease phenotype, prognosis, and therapy response are also affected by the temporal order of these lesions [70] and progenitor stage where they appear [71], recently studied at single-cell resolution [72,73]. This heterogeneity is also apparent in single-cell gene expression [16,17]. The dynamics of chromatin landscape in clonal evolution have not been studied to the same extent, despite strong evidence of the essential role of chromatin modifiers.

Here we analyzed genome-wide dynamics of transcription and chromatin marks during clonal evolution and showed how these dynamics reinforce a core cellular program enabling growth advantage. Despite various caveats of mouse MLL-AF9 overexpressing cells as a model of human disease, this basic but tractable model enabled us to make predictions that were later confirmed by AML patient data. Methodologically, our muliomic analyses of clonal evolution in independent leukemic clones proved an effective new approach to identify candidate genes for common causal regulators of clonal growth. In fact, the gradual nature of gene regulation during clonal evolution would make it challenging to detect some of these key regulators using traditional screening approaches such as CRISPR screens.

Unexpectedly, most of the promoters and enhancers affected by clonal evolution had been already affected at the early leukemic stage of the same clone (Figures 2H, 3F, S2L, S3H). Even more surprisingly, rather than being further amplified, these early changes were preferentially reversed during clonal evolution (Figures 2I, 3G, S2M, S3I). This reversal is consistent with the reported changes in the roles of some central leukemic regulators at later stages of AML. Polycomb component Ezh2 is a strong tumor suppressor during AML induction and is later switched to an oncogenic role during AML maintenance [14], and Igf1r is a major oncogene during AML induction but is not required during later clonal growth [52]. The biological role of this reversal is an important question. As one possible explanation, the initial leukemogenic lesion (MLL rearrangement) causes perturbations across a wide variety of direct and indirect genomic targets (Figure 1G,L), some of which (activation of Hoxa9, Meis1, etc) cause oncogenic transformation, while others may be non-essential and even maladaptive for later clonal fitness and growth. In this case, scaling back maladaptive initial effects (Figure 2C, 3C, 4D,F) while keeping a strong initial activation of leukemogenic drivers (Figure 1F) may provide a growth advantage. This reversal may be achieved through the regulation of promoters and enhancers by transcription factors and chromatin regulators.

Our further goal was to identify a gene set that would be enriched in causal regulators of leukemic cell growth shared between evolving clones. For that, we leveraged the variability of evolutionary paths in individual clones and stringently filtered all genes for a full consistency among the changes of their expression and chromatin marks in both clones in vivo and ex vivo (Figure 4A). This filtering revealed a small set of core genes (Figure 4B). We selected three of these genes for experimental validation and confirmed their causal roles as two leukemic cell growth suppressors (Irx5 and Plag1) and one driver (Smad1). Irx5 is a curious example of a strong MLL-AF9 target [49] that was released from Polycomb repression and dramatically activated at the early leukemic stage, like known major leukemogenic drivers (Figure 4D). However, unlike these drivers, during clonal evolution Irx5 reversed this activation by increased Polycomb repression (Figure 4D). Plag1 is an oncogene derepressed at the induction of AML [14,50], but its surprising role in suppressing cell growth at the later stages of clonal evolution has not been previously reported. SMAD1 has not been reported as a major leukemic regulator, although it is a central effector of the BMP pathway implicated in leukemia [38,39]. Smad1 was the only Smad family member with this behavior, which positions BMP pathway as a possible therapeutic target distinct from the related TGFβ pathway.

Downstream transcriptional effects of these three genes upon their overexpression converge on a network of transcription factors, chromatin modifiers, and components of DNA methylation machinery, many strongly implicated in leukemia (Figure 6I). The effects on Smad3, a key effector of TGFβ pathway (Figure 6I), may suggest that Smad3-mediated TGFβ signaling is inhibited and replaced by Smad1-mediated BMP signaling during clonal evolution. This inhibition is in line with the observed regulation of other TGFβ related genes (Peg10, Matn2, Ptpn3, Figure 4B). Coordinated downstream regulation by Irx5, Plag1, and Smad1 also affected major epigenetic factors: H3K4me2/3 demethylase Kdm5b, Polycomb component Bcorl1 [58,59], as well as DNA methylase Dnmt3a [7] and DNA demethylation component Tet2 [8], highlighting the role of repressive Polycomb, H3K4 demethylation, and DNA methylation machineries. Modulation of Polycomb repression was a major mechanism of promoter regulation at both the early leukemic stage and during the later clonal evolution, for example at *Smad1*, *Irx5*, and *Plag1* promoters (Figure 4C-F). The reported role of Kdm5b as aggressiveness suppressor in leukemia [15] was consistent with its transcriptional regulation by Smad1, Irx5, and Plag1 (Figure 6E). DNA methylation and demethylation components with known strong AML involvement[7,8] were regulated by Smad1 and Plag1 in opposite directions (Figure 6E). This collective involvement of epigenetic repressors is consistent with their known functional crosstalk. In addition to antagonism between Polycomb and H3K4 methylation [23], as well as direct Polycomb repression of Kdm5b [15], there is growing evidence of direct Polycomb-DNMT3A interactions [74] contributing to leukemia [75].

Importantly, our predictions derived from MLL-AF9 overexpressing mouse cell model were later confirmed by the gene expression and survival data in AML patients. Consistent with our predictions, IRX5 is upregulated across a wide range of AML subtypes compared to normal controls, while SMAD1 is downregulated specifically in MLLr leukemias (Figure 6F). Furthermore, higher IRX5 and SMAD1 expression levels corresponded to longer (Figure 6I, top) and shorter (Figure 6I, bottom) patient survival, respectively. This suggested the value of mutiomic analyses of clonal evolution in a mouse cell model as a predictive method applicable to human leukemia.

Taken together, our results suggest that evolution of leukemic clones is often mediated through gradual adjustment of chromatin states and gene expression. This adjustment often involved a surprising reversal of prior changes that had occurred at the early leukemic stage. Through genome-wide analyses of chromatin and gene expression dynamics, we identified a set of core genes whose changes during clonal evolution were shared between individual clones and cell environments, and successfully validated three of these genes as causal regulators of leukemic cell growth. Two of these genes, were further confirmed as markers of AML status in human patient data.

## MATERIALS AND METHODS

### Murine models and cell lines

Experiments were approved by the Institutional Animal Care and Use Committee (IACUC) of the Massachusetts General Hospital. C57BL/6J, B6.SJL-Ptprca Pepcb/BoyJ (CD45.1) were purchased from Jackson Laboratories. All MLL-AF9 leukemic cell lines were obtained as previously described [76] and maintained in RPMI1640 (10% FBS), penicillin/streptomycin, 6 ng/ml rmIL-3, and approximately 100 ng/ml SCF. Briefly, a MLL-AF9 CD45.1 parental line was established by sorting, using a FACS AriaII (BD Biosciences), granulocyte-macrophage progenitors (GMP; c-Kit^high^Sca-1^−^ Gr-1^low^CD16/32^+^CD34^+^) from a male CD45.1 donor. Retrovirally transfection with a MSCV-MLL/AF9-IRES-NeoR construct in sorted progenitor cells was performed a day after incubation in RPMI1640 (Lonza) transduction medium containing 10% (FBS; Gibco), 100 I.U./ml penicillin, 100 mg/ml streptomycin (Gibco), 10 ng/ml recombinant mouse SCF (rmSCF), 6 ng/ml recombinant mouse interleukin 3 (rmIL-3), 5 ng/ml rmIL-6 (all from Peprotech). Progenitors were transferred to RetroNectin®-coated (Lonza) 6-well plates in a volume of 1 ml, combined with 1 ml of fresh retroviral supernatant packaged in 293FT cells (Invitrogen) with pCL-Eco (Addgene 12371) and 8 µg/ml final concentration of polybrene, then spinoculating (1,000g, 90 min, 22°). Immediately following the transduction, 4 ml of fresh transduction medium was added to each well to dilute the polybrene and 24 hours post-transduction, cells were washed by centrifugation (500g, 5 min, 4°C) then resuspended in fresh transduction medium. 48 hours post-transduction, neomycin resistant lines were selected using 50 µg/ml neomycin (Gibco).

### Clonal fluorescent labeling

To generate MLL-AF9 clones expressing fluorescent proteins (FP), single MLL-AF9 CD45.1 clones were sorted using a FACS Aria II (BD Biosciences) into 96-well plates. A single clone was expanded and transduced with lentiviral constructs EF1a-FP-SV40-PuroR, generated by Gateway cloning of FP Entry plasmids into the pLEX_307 Destination construct (a gift from David Root, Addgene plasmid # 41392). From each FP transduction, single subclones were expanded in vitro and those that stably expressed FPs were preserved.

### Syngeneic leukemia experiments

All leukemia transplantations were performed by injecting cells intravenously (tail vein) in a volume of 300 ul of phosphate-buffered saline (PBS) in recipient C57Bl/6J male mice (age 8-10 weeks) that were sublethally irradiated (4.5Gy) the day prior using a Cesium-137 source. Blood was collected through tail vein sampling and complete blood counts were done on a Heska Element HT5 analyzer. Bone marrow aspiration was done under isoflurane anaesthesia using a 25G needle attached to a 1 ml syringe rinsed with PBS, gently inserted into the femoral cavity through the articular surface of the knee; approximately 10 µl of bone marrow was gently aspirated in the syringe and immediately resuspended in 300 µl of PBS on ice. Hematopoietic cell isolation was done as follows: after euthanasia by asphyxiation with CO_2_, vertebrae and the long bones of the arms and legs were dissected and crushed into PBS containing 2% FBS (FACS buffer). The bone marrow mononuclear cells were isolated over a Ficoll-Paque PLUS density gradient and counted using trypan blue and the Cellometer instrument.

### Overexpression of gene candidates

Leukemic subclones were cultured in RPMI 1640 medium supplemented with 10% fetal bovine serum, 1% L-glutamine, 1% penicillin/streptomycin, 25 ng/mL SCF, and 5 ng/mL IL-3. ORFs of interest were overexpressed using second-generation lentiviral system that consisted of pCMV-VSV-G (Weinberg lab) envelope plasmid and PsPAX2 (Trono lab) packaging plasmid acquired from Addgene. HEK293t cells were transfected with plasmid DNA at a ratio of 5:4:1 transfer plasmid:envelope plasmid:packaging plasmid. Transfection efficiency was aided by the addition of Promega’s Fugene6 Reagent at a ratio of 3:1 to plasmid DNA. HEK293t cells were cultured in DMEM, high glucose, pyruvate supplemented with 10% fetal bovine serum and 1% penicillin/streptomycin. Lentiviral titer was assessed by FACS. Mouse leukemic progenitors were transduced by exposing them to viral particles and room temperature centrifugation at 1000g for 1 hr in 8uG/mL polybrene (Millipore Sigma). Spinfection plates were pre-coated using 500 uL of PBS with 10 uG/mL of purified human plasma fibronectin (Sigma-Aldrich) and incubated at 37°C several hours prior to transduction. Single cell FACS sorting was performed 48 hours after transduction to isolate individual GFP-positive clones. Overexpression of the gene of interest in selected clones was confirmed by RT-qPCR and RNA-seq. RNA was extracted using TRIzol LS Reagent, followed by cDNA synthesis using SuperScript First-Strand Synthesis System for RT-PCR (Invitrogen).

### Cell growth quantitation

Cell growth was quantitated using CellTrace Far Red Cell Proliferation Kit (Invitrogen), Initially, on day 1, cell lines were standardized to a density of 1,000,000 cells/mL in 1mL of PBS, labeled with CellTrace™ Far Red dye at a 1 μM concentration, and subsequently centrifuged at 500g for 5 minutes and resuspended in fresh media. The labeled cells were then cultured in 6-well plates containing 5 mL of RPMI 1640 media supplemented with 10% Fetal Bovine Serum. Starting on day 2, daily sampling was conducted by aspirating a fraction of the culture, centrifuging at 500g for 5 minutes, and resuspending the cell pellet in 400 µL FACS Buffer containing 0.8% formaldehyde for overnight fixation. The following day, formaldehyde fixation was neutralized with the addition of 100 µL FBS. These fixed samples were stored at 4°C until the final timepoint. Flow cytometric analysis was performed on a BD FACS Melody system, where the mean fluorescence intensity of each sample was measured at every timepoint. The doubling time was inferred by calculating the time interval required for the fluorescence intensity to halve, reflecting cell proliferation rates.

### Cell sorting to assess hematopoietic lineage

Cellular suspensions were prepared for flow cytometric analysis by harvesting 1 million cells and resuspending them in 100 µL of FACS Buffer, consisting of 2% fetal bovine serum, 1 mM EDTA, and phosphate-buffered saline. The cell suspensions were then labeled with lineage related antibodies: PE/Dazzle™ 594 anti-mouse/human CD11b (Biolegend 101256), APC anti-mouse Ly-6G/Ly-6C (Gr-1) (Biolegend 108412), PE anti-mouse CD123 (Biolegend 106005), and PE/Cyanine7 anti-mouse CD117 (c-kit) (Biolegend 135112), at the appropriate concentrations for the volume and cell count (1uL in 1 million cells), followed by incubation at room temperature for 30 minutes in the dark. Post-incubation, cells were centrifuged at 500g for 5 minutes, washed twice with FACS Buffer, and finally resuspended in 400 µL of the same buffer. The cell suspensions were then strained through a 70 µm mesh into FACS tubes to ensure a single-cell suspension. Single-color controls were established using OneComp eBeads™ (Thermofisher 01-1111-41). Analysis was performed in FlowJo, with doublet discrimination and fluorescent compensation.

### Cell Cycle Assay

Cells were grown out for two days and resuspended at 10 million cells/mL in 1X Dulbecco’s Phosphate Buffered Saline (DPBS) (Life Technologies Limited, Paisley, PA, USA). 3 million cells were then transferred to a FACS tube and chilled on ice for 5 minutes. Ice cold ethanol (Sigma-Aldrich, Burlington, VT, USA) was added while samples were vortexed. Tubes were then stored at -20_°_C overnight. 3mL of 1X DPBS was added to each tube and tubes were spun down at 500g for 5 minutes. Cells were then resuspended in 300µl of staining buffer (0.4 mg/mL RNAse A (Thermofisher, Waltham, MA, USA), 10 uG/mL propidium iodide (Life Technologies Corporation, Eugene, OR, USA), 0.1% Triton X-100 (Sigma-Aldrich, St. Louis, MO, USA) in PBS) and filtered through a 40µm Corning Cell Strainer (Corning Incorporated, Durham, NC, USA) into FACS tubes. Samples were then incubated at 37_°_C for 15 minutes and flowed on a Becton-Dickinson (BD) FACSCelesta instrument to collect 10,000 events per sample. Flow cytometry data was analyzed using FlowJo v10.10 (BD Biosciences, Ashland, OR, USA).

### Annexin V Apoptosis Assay

A modified protocol provided by Biotium was used to prepare samples. 5X Annexin V Binding Buffer (Biotium, Fremont, CA, USA) was first diluted with Ambion nuclease-free water (Invitrogen, Austin, TX, USA) at a 1:5 ratio. Cultured cells were washed with 1X Dulbecco’s Phosphate Buffered Saline (DPBS) (Life Technologies Limited, Paisley, PA, USA) once and resuspended at 3,000,000 cells/mL in diluted binding buffer. Annexin V Conjugate CF 633 (Biotium, Fremont, CA) was added to 100µl of each cell type for a final concentration of 1µg/mL. Cells were incubated at room temperature for 30 minutes in the dark and then washed with 1X DPBS once before staining with 4’,6-diamidino-2-phenylindole, dihydrochloride (DAPI) (Invitrogen, Austin, TX, USA). 1X Binding Buffer was then added to each tube and tubes were flowed on a Becton-Dickinson (BD) FACSCelesta instrument to collect 10,000 events per cell line. Flow cytometry data was analyzed using FlowJo v10.10 (BD Biosciences, Ashland, OR, USA).

### Methylcellulose colony forming assay

A modified protocol from StemCell Technologies was used to culture cell lines in methylcellulose. MethoCult GF M3434 (StemCell Technologies, Canada) was thawed overnight at 4_°_C. Cells were counted using Acridine Orange/Propidium Iodide Stain (Aligned Genetics, Inc., Annandale, VA, USA) and diluted to 50,000 cells/mL. Cells were counted again and ∼1500 cells were added to each MethoCult GF M3434 tube. Tubes were shaken vigorously and allowed to settle for 5 minutes. A 3mL syringe and 18 gauge 1-1/2 inch needle were used to plate 1.1mL of MethoCult in 35mm plates in duplicate. Each 35mm plate was placed inside a 10cm plate along with another 35mm plate filled with sterile water. All plates were incubated at 37_°_C, 5% CO_2_ and counted after eight days. Mean and standard deviation based on biological duplicates were calculated for each cell type.

### Assaying effects of BMP inhibitor on cell growth ex vivo

ER-Hoxb8 overexpressing progenitors used as normal control were cultured in RPMI1640 (10% FBS), 1% penicillin/streptomycin, 1% L-glutamine, 100 ng/ml SCF and 0.5 uM β-estradiol. Early and aggressive leukemic subclones as well as normal ER-Hoxb8 overexpressing progenitors were grown out, resuspended at 50,000 cells/mL in their respective media, and plated in 96-well plates at 100µl of cells per well. Cells were then treated with 10µM to 0.005µM concentrations of inhibitor LDN-212854 (Selleck Chem, Houston, TX, USA) using the D300e Digital Dispenser (Tecan Trading AG, Switzerland). Plates were then incubated at 37_°_C for 3 days, after which 100µl of diluted CellTiter-Glo 2.0 Cell Viability Assay (Promega, Madison, WI, USA) (1:1 in 1X Dulbecco’s Phosphate Buffered Saline (DPBS) (Life Technologies Limited, Paisley, PA, USA) was applied to each well and mixed thoroughly. Plates were then read using a BioTek Synergy HTX instrument (BioSPX, Netherlands) for fluorescence and then analyzed on Prism 10.0 for IC50 (GraphPad, Boston, MA, USA).

### RNA library preparation and sequencing (RNA-seq)

Total RNA was isolated from three biological replicates of each cell type using TRIzol (Invitrogen) and DNase treated. Sequencing libraries were prepared using polyA-selection followed by NEBNext Ultra Directional RNA library preparation protocol (New England Biolabs). Sequencing was performed on Illumina HiSeq 2500 instrument, resulting in approximately 30 million 50 bp reads per sample.

### Chromatin immunoprecipitation library preparation and sequencing (ChIP-seq)

Cells were crosslinked for 10 minutes in 1% formaldehyde at room temperature and quenched with 125mM Glycine. ChIP was performed as previously described [77]. Antibodies used: H3K4me1 (Abcam ab8895), H3K4me3 (Millipore, 04-745), H3K27me3 (Cell Signaling Technology, C36B11), H3K27ac (Abcam ab4729). Samples were further processed for high throughput sequencing. Sequencing was performed on Illumina HiSeq 2500 instrument resulting in approximately 30 million paired-end 50 bp reads per sample.

### RNA-seq data analyses

Sequencing reads were mapped to the mouse transcriptome (mm9 EMSEMBL annotation) using STAR [78]. Gene expression counts were calculated using HTSeq v.0.6.0 [79]. Calculation of expression values and differential expression analysis were performed using EgdeR package [80] after normalizing read counts and including only genes with count per million reads (CPM) > 1 for one or more samples. Differentially expressed genes were defined based on the criteria of > 2 fold change of expression and FDR < 0.05. Analysis of gene set enrichment was performed using Enrichr [22].

### ChIP-seq data analyses

Reads were aligned against the mm9 reference genome using BWA [81]. Alignments were filtered for uniquely mapped reads and alignment duplicates were removed. Input-normalized and CPM normalized coverage tracks were generated using the bamcompare function of deepTools [82]. Enrichment over TSS-proximal regions (TSS ± 3 Kb) was calculated as the ratio of ChIP and input read counts scaled by library size. Reads of publicly available MLL-AF9 ChIP-seq (GEO accession number GSE29130) were aligned as described previously, and peaks were identified using Homer with the “factor” option. Strong targets genes of MLL-AF9 were identified by filtering the peak list using a peak enrichment cutoff >=13. Putative enhancer elements were identified using H3K27ac and H3K4me1 ChIP-Seq peaks. In brief, we analyzed tag counts in sliding 1 Kb windows with the step of 200 bp and estimated statistical significance of enrichment of ChIP vs input using negative binomial distribution with the parameter of mean based on input tag count and the size parameter (s) selected based on manual inspection of peak calls. Regions of significant enrichment were generated by merging adjacent significantly enriched windows within 1 Kb. To define putative enhancer elements, peak sets of H3K27ac and H3K4me1 were merged, followed by removing regions within 3 Kb from annotated promoters and regions overlapping H3K4me3 peaks.

### Analyses of public human single-cell RNA-seq data

Publicly available scRNA-seq read counts from GEO dataset GSE116256 [16] were analyzed using Seurat 3.2.3 package [83]. We used the patient samples with > 300 cells annotated as GMP or GMP-like by the authors. Cells with <200 expressed genes and genes expressed in <3 cells were filtered out, followed by the exclusion of cells with high content of mitochondrial transcripts (> 10% of total reads). Counts across all cells for each sample were normalized using NormalizeData function in Seurat. Individual sub-samples were integrated using Seurat canonical correlation analysis (CCA) method. Integration anchors were determined using FindIntegrationAnchors function and then used in IntegrateData function. We used cell annotations provided in the study to visualize expression of genes of interest in GMP and GMP-like cells using DotPlot function of Seurat.

## DECLARATIONS

### Ethics approval

Not applicable

### Consent for publication

Not applicable

### Availability of data and materials

The datasets generated and analyzed during the current study are available in the GEO repository with accession numbers GSE280194 for RNA-seq and GSE280195 for ChIP-seq data.

### Author contributions

RIS, FM, DSK, DS, and REK contributed to conception and design of the work. AM, SK, AD, GK; NEJ, EEY, JM, TZ, JX, RB, and MM contributed to the acquisition and interpretation of experimental data. GB and KC performed computational data analyses and interpretation. GB, RIS, and DS have drafted the manuscript. RIS, DS, FM, and NEJ substantively revised it. All authors reviewed the manuscript.

