## Supplemental Figures for "Multiomic analysis of clonal development reveals new regulators of leukemic cell growth"

**Bonilla et al.**

### Supplementary Figures

Figure S1

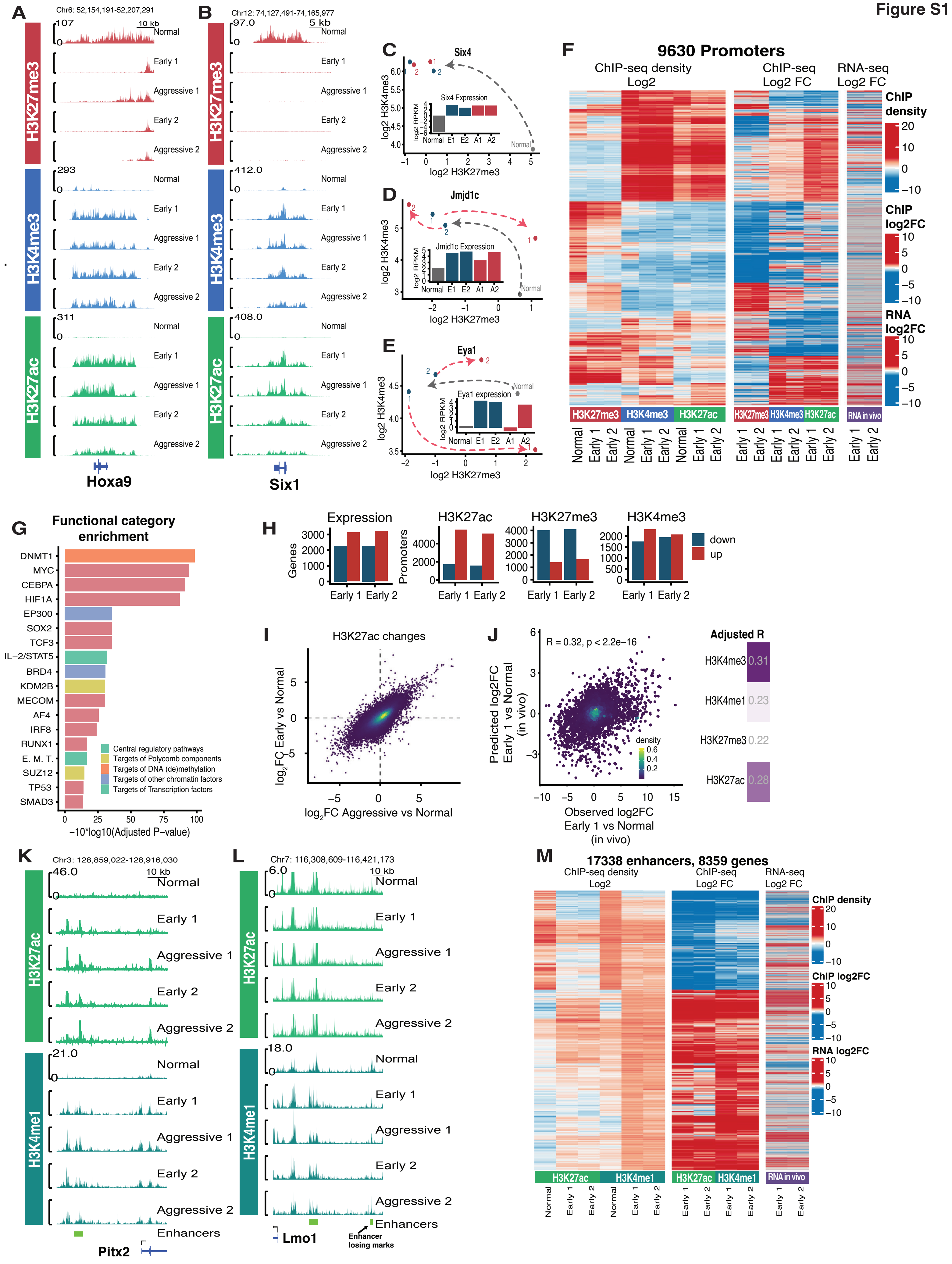
A
Chr6: 52,154,191-52,207,291
10 kb
Normal
Early 1
Aggressive 1
Early 2
Aggressive 2
H3K27me3
H3K4me3
H3K27ac
Hoxa9
B
Chr12: 74,127,491-74,165,977
5 kb
Six1
C
Six4
Six4 Expression
log2 H3K4me3
log2 H3K27me3
log2 RPKM
Normal E1 E2 A1 A2
D
Jmjd1c
Jmjd1c Expression
E
Eya1
Eya1 expression
F
9630 Promoters
ChIP-seq density Log2
ChIP-seq Log2 FC
RNA-seq Log2 FC
ChIP density
ChIP log2FC
RNA log2FC
RNA in vivo
G
Functional category enrichment
DNMT1
MYC
CEBPA
HIF1A
EP300
SOX2
TCF3
IL−2/STAT5
BRD4
KDM2B
MECOM
AF4
IRF8
RUNX1
E. M. T.
SUZ12
TP53
SMAD3
Central regulatory pathways
Targets of Polycomb components
Targets of DNA (de)methylation
Targets of other chromatin factors
Targets of Transcription factors
−10*log10(Adjusted P−value)
H
Expression
H3K27ac
H3K27me3
H3K4me3
Genes
Promoters
Early 1 Early 2
down
up
I
H3K27ac changes
log2FC Early vs Normal
log2FC Aggressive vs Normal
J
R = 0.32, p < 2.2e−16
Predicted log2FC Early 1 vs Normal (in vivo)
Observed log2FC Early 1 vs Normal (in vivo)
density
Adjusted R
H3K4me3 0.31
H3K4me1 0.23
H3K27me3 0.22
H3K27ac 0.28
K
Chr3: 128,859,022-128,916,030
10 kb
H3K27ac
H3K4me1
Enhancers
Pitx2
L
Chr7: 116,308,609-116,421,173
10 kb
Enhancers
Lmo1
Enhancer losing marks
M
17338 enhancers, 8359 genes
ChIP-seq density Log2
ChIP-seq Log2 FC
RNA-seq Log2 FC
ChIP density
ChIP log2FC
RNA log2FC

**Figure S1 (related to Figure 1). At the early leukemic stage, changes of promoter and enhancer chromatin state and gene expression across the genome were consistent between independent clones and were mostly maintained at the later aggressive stage. A-B.** Additional examples of ChIP-seq genomic tracks for known major drivers of MLLr leukemia: Hoxa9 and Six1. Input-normalized ChIP-seq density of H3K27me3 (red), H3K4me3 (blue), and H3K27ac (green) in normal progenitors (top) compared to two independent leukemic clones 1 and 2 at different stages of aggressiveness (early and aggressive). Promoter chromatin state was dramatically activated in all leukemic subclones compared to normal but showed no strong consistent difference between two aggressiveness stages. **C-E.** Chromatin state progression in the space of H3K27me3 vs H3K4me3 for promoters of known major drivers of MLLr leukemia: Six4, Jmjd1c, Eya1. Normal progenitors (grey) and early (blue) and aggressive (red) stages of two leukemic clones (1 and 2) are represented as points in the space of TSS-proximal H3K27me3 (x axis) vs H3K4me3 density (y axis). The corresponding gene expression levels (log2 RNA-seq RPKM) are shown as barplots. The early loss of repressive H3K27me3 mark accompanied by a strong gain of active H3K4me3 mark and upregulation of expression was later maintained at the aggressive stage. **F.** Expanded heatmap of early changes (log2 fold change) of promoter chromatin mark densities across all promoters with a substantive level and > 2-fold change of at least one mark (middle), juxtaposed with the absolute levels of mark densities in normal progenitors and two clones at early and aggressive stage (left) and changes of gene expression in vivo (log2 fold change of RPKM, right). Both absolute mark levels and their changes were strongly consistent between two independent clones (two columns for each mark and expression). **G.** Functional category enrichment among genes differentially expressed between normal and early leukemic progenitors in vivo consistently in both clones. Bars colored by category types represent adjusted P-value of enrichment according to EnrichR. **H.** In both clones, early-stage changes were skewed towards the increase of active H3K27ac and H3K4me3 promoter marks and gene expression and decrease of repressive mark H3K27me3. Barplots of promoter numbers with increase (red) and decrease (blue) of each individual mark and gene expression at the early stage in each leukemic clone (Early 1, Early 2). **I.** Most of the early changes of promoter chromatin marks and expression were later maintained at the same level during clonal evolution. Scatterplot of H3K27ac changes (log2 fold change) at all promoters between normal cells and the early leukemic stage (y axis) compared to the changes between normal cells and the aggressive leukemic stage (x axis). Individual promoters are represented as points, color indicates point density. **J.** Combined changes of promoter chromatin marks were predictive of changes of gene expression. Scatterplot of observed change of gene expression at the early stage of clone 1 (log2 of fold change, x axis) vs predicted change (y axis) based on the combination of changes (log2 fold change) of promoter H3K27ac, H3K4me1, H3K4me3, and H3K27me3. The accuracy of predictions based on individual single marks is shown as a heatmap on the right. **K-L.** Additional examples of ChIP-seq genomic tracks at enhancers that changed chromatin state at the early leukemic stage. Input-normalized ChIP-seq densities of H3K27ac (green) and H3K4me1 (dark green) in normal progenitors (top) compared to two independent leukemic clones 1 and 2 at two stages of aggressiveness. **M.** Expanded heatmap of early changes (log2 fold change) of H3K27ac and H3K4me1 densities across all enhancers with a substantive level and > 2-fold change of at least one mark (middle), juxtaposed with the absolute levels of mark densities in normal progenitors and two leukemic clones at early and aggressive stage (left) and changes of proximal gene expression in vivo (log2 fold change of RPKM, right). Both absolute mark levels and their changes were strongly consistent between two independent clones (two columns for each mark and expression)

Figure S2

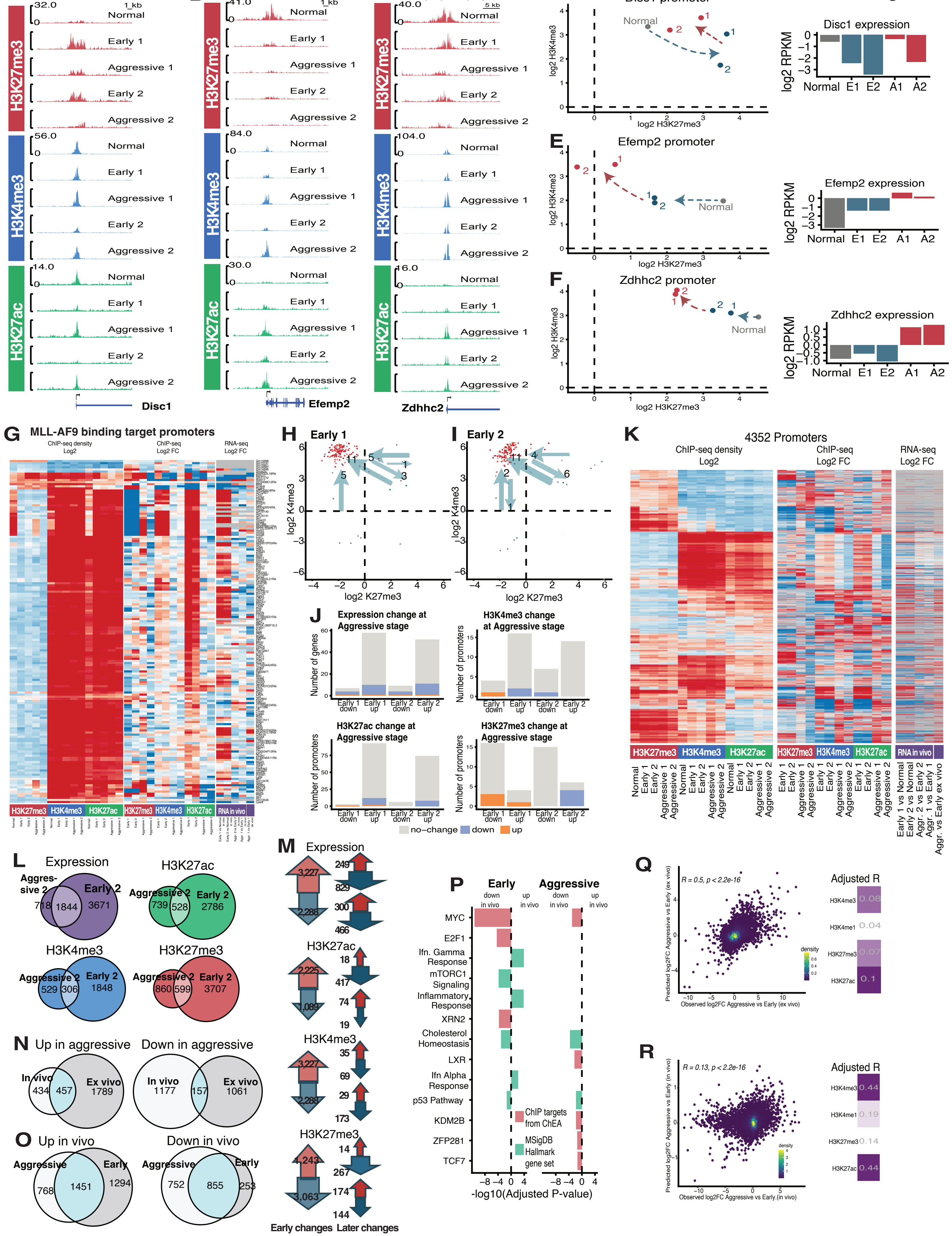
A
Chr8: 127,570,043-127,589,776
1 kb
Normal
Early 1
Aggressive 1
Early 2
Aggressive 2
H3K27me3
H3K4me3
H3K27ac
Disc1
B
Chr19: 5,467,196-5,486,707
Efemp2
C
Chr8: 41,495,822-41,525,133
5 kb
Zdhhc2
D
Disc1 promoter
log2 H3K4me3
log2 H3K27me3
Disc1 expression
log2 RPKM
Normal
E1
E2
A1
A2
E
Efemp2 promoter
Efemp2 expression
F
Zdhhc2 promoter
Zdhhc2 expression
G
MLL-AF9 binding target promoters
ChIP-seq density Log2
ChIP-seq Log2 FC
RNA-seq Log2 FC
RNA in vivo
H
Early 1
log2 K4me3
log2 K27me3
I
Early 2
J
Expression change at Aggressive stage
Number of genes
H3K4me3 change at Aggressive stage
Number of promoters
H3K27ac change at Aggressive stage
H3K27me3 change at Aggressive stage
Early 1 down
Early 1 up
Early 2 down
Early 2 up
no−change
down
up
K
4352 Promoters
ChIP-seq density Log2
ChIP-seq Log2 FC
RNA-seq Log2 FC
Early 1 vs Normal
Early 2 vs Normal
Aggr. 2 vs Early 2
Aggr. 1 vs Early 1
Aggr. vs Early ex vivo
L
Expression
Aggressive 2
Early 2
718
1844
3671
H3K27ac
739
528
2786
H3K4me3
529
306
1848
H3K27me3
860
599
3707
M
Expression
3,227
2,288
249
829
300
466
H3K27ac
2,225
1,089
18
417
74
19
H3K4me3
3,227
2,288
35
69
29
173
H3K27me3
4,243
3,063
14
267
174
144
Early changes
Later changes
N
Up in aggressive
Down in aggressive
In vivo
Ex vivo
434
457
1789
1177
157
1061
O
Up in vivo
Down in vivo
Aggressive
Early
768
1451
1294
752
855
253
P
Early
Aggressive
down in vivo
up in vivo
MYC
E2F1
Ifn. Gamma Response
mTORC1 Signaling
Inflammatory Response
XRN2
Cholesterol Homeostasis
LXR
Ifn Alpha Response
p53 Pathway
KDM2B
ZFP281
TCF7
ChIP targets from ChEA
MSigDB Hallmark gene set
-log10(Adjusted P-value)
Q
R = 0.5, p < 2.2e-16
Predicted log2FC Aggressive vs Early (ex vivo)
Observed log2FC Aggressive vs Early (ex vivo)
density
Adjusted R
H3K4me3 0.08
H3K4me1 0.04
H3K27me3 0.07
H3K27ac 0.1
R
R = 0.13, p < 2.2e-16
Predicted log2FC Aggressive vs Early (in vivo)
Observed log2FC Aggressive vs Early (in vivo)
Adjusted R
H3K4me3 0.44
H3K4me1 0.19
H3K27me3 0.14
H3K27ac 0.44

**Figure S2 (related to Figure 2). Changes of promoter chromatin states and expression at the aggressive stage were more clone-specific, gradual, and often reversed early leukemic changes. A-C.** Additional examples of ChIP-seq genomic tracks showing promoter chromatin mark and expression changes between two stages of aggressiveness. H3K27me3 (red), H3K4me3 (blue), and H3K27ac (green) ChIP-seq genomic tracks in normal progenitors (top) compared to two independent leukemic clones 1 and 2 at two stages of aggressiveness. Tracks shown for promoters of Disc1 **(A)**, Efemp2 **(B)** and Zdhhc2 **(C)**. **D-F.** Chromatin state progression in the space of H3K27me3 vs H3K4me3 for promoters shown in Figure S2 A-C. Normal progenitors (grey) and early (blue) and aggressive (red) stages of two leukemic clones (1 and 2) are represented as points in the space of H3K27me3 (x axis) vs H3K4me3 density (y axis) in the TSS-proximal region. The corresponding gene expression levels (RNA-seq RPKM) are shown as barplots. **G.** In contrast to the early stage, the vast majority of strong MLL-AF9 targets did not change their chromatin state or expression at the aggressive stage. Heatmap of chromatin mark and gene expression progression for the strongest MLL-AF9 targets according to previously published ChIP-seq data (39). Left, absolute values of promoter chromatin mark densities. Second left, log2 fold change of chromatin marks between stages. Right, log2 fold change of expression between stages. **H-J.** Chromatin states of the strongest MLL-AF9 targets and their changes at the early stage, shown as scatterplots in the space of H3K27me3 vs H3K4me3 densities. Strong skew towards activation is shown by the arrows with the corresponding promoter numbers for each direction. **H-I.** Changes at the early stage in two leukemic clones (1 and 2). **J.** Barplots showing proportions of promoters of strongest MLL-AF9 targets increasing (orange), decreasing (blue), or not changing (fold change < 2) their expression (top left) and chromatin marks among the groups of promoters that showed increase or decrease at the early stage. **K.** Expanded heatmap of changes (log2 fold change) of promoter chromatin mark densities between early and aggressive leukemic stage in two clones across all promoters with a substantive level and > 2-fold change of at least one mark, juxtaposed with the absolute levels of mark densities in normal progenitors and two leukemic clones at early and aggressive stage (left) and the changes of gene expression in vivo (log2 fold change of RPKM, right). Absolute levels and changes at the aggressive stage were only partially consistent between two independent clones (two columns for each mark and expression). **L.** Most promoters affected at the aggressive stage underwent prior changes at the early leukemic stage. Venn diagrams of promoters with changes of expression and individual chromatin marks at the aggressive and early stages for clone 2. **M.** Clonal evolution preferentially reversed the prior early leukemic changes at the promoters. Large arrows: numbers of promoters with increase (red) and decrease (blue) of expression and chromatin marks (H3K27ac, H3K4me3, H3K27me3) between normal progenitors and the early leukemic subclone. Smaller arrows near each large arrow: numbers of these promoters that underwent the later increase (red) or decrease (blue) of expression or chromatin mark during clonal evolution. These later changes were skewed in the direction opposite to the early leukemic changes. Data for clone 2. **N.** Overlap of changes between aggressive stage and earlier leukemic stage in vivo compared to ex vivo. **O.** DEGs associated with the difference between in vivo and ex vivo conditions were strongly overlapping but still different between the early and aggressive subclones. Venn diagrams of up- and downregulated genes in vivo vs ex vivo for the early and aggressive subclones. **P.** Overlap of functional categories enriched among genes up- and downregulated (bars to the right and left of dotted line, respectively) in vivo vs ex vivo for the early and aggressive leukemic subclones. Bars colored by category types represent log10 of adjusted P-value of enrichment according to EnrichR. **Q.** Combined changes of promoter chromatin marks were predictive of changes of gene expression ex vivo. Scatterplots of observed change of gene expression at the aggressive vs early stage of clone 1 (log2 of fold change, x axis) plotted against predicted change (y axis) based on the combination of changes (log2 fold change) of promoter H3K27ac, H3K4me1, H3K4me3, and H3K27me3. The accuracy of predictions based on individual single marks is shown as a heatmap on the right of each scatterplot. **R.** Combined changes of promoter chromatin marks were predictive of changes of gene expression in vivo (similar to **Q**).

**Figure S3**

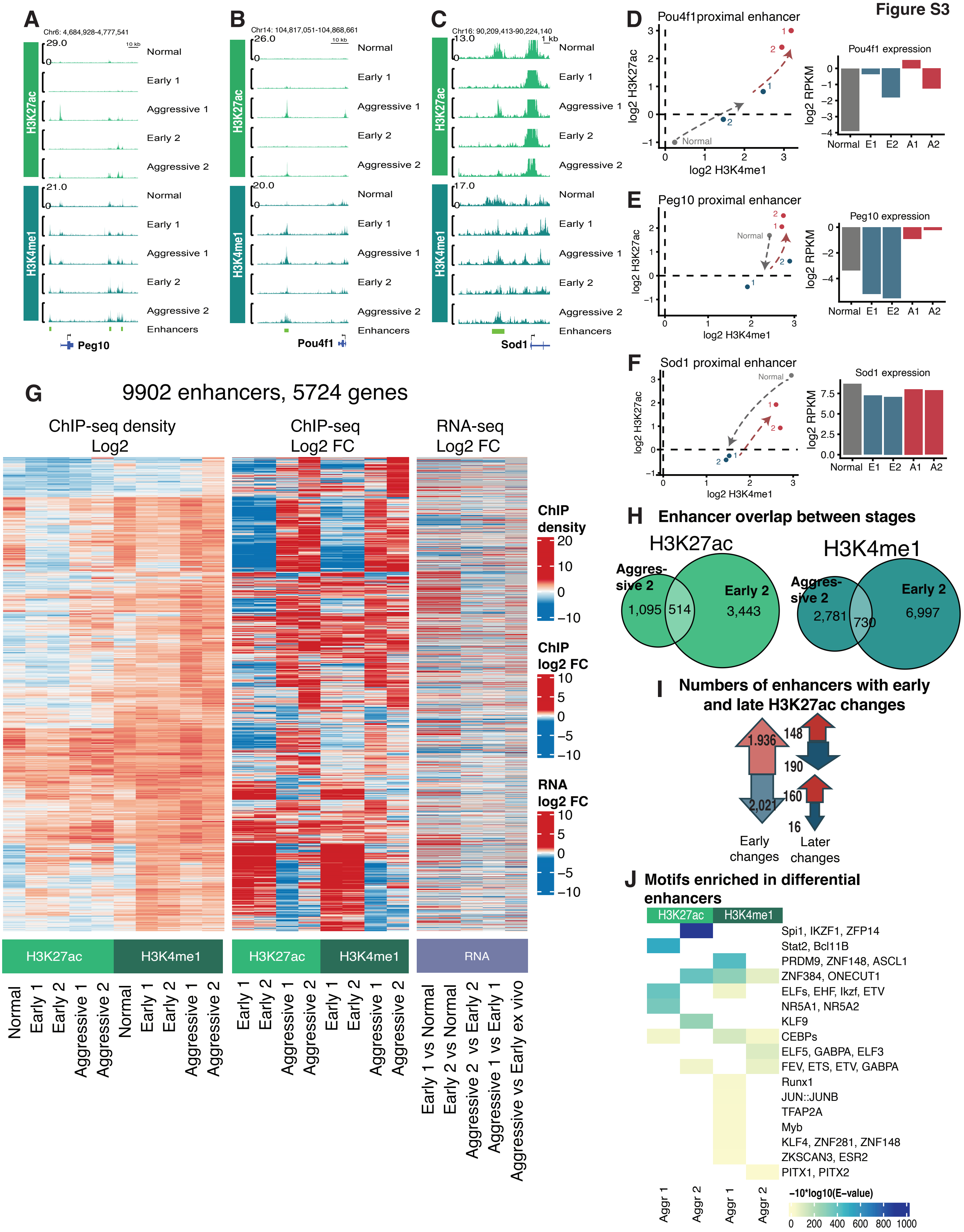
A
Chr6: 4,684,928-4,777,541
29.0
10 kb
Normal
Early 1
Aggressive 1
Early 2
Aggressive 2
21.0
Enhancers
Peg10
H3K27ac
H3K4me1
B
Chr14: 104,817,051-104,868,661
26.0
20.0
Pou4f1
C
Chr16: 90,209,413-90,224,140
13.0
17.0
1 kb
Sod1
D
Pou4f1proximal enhancer
log2 H3K27ac
log2 H3K4me1
Pou4f1 expression
log2 RPKM
Normal E1 E2 A1 A2
E
Peg10 proximal enhancer
Peg10 expression
F
Sod1 proximal enhancer
Sod1 expression
G
9902 enhancers, 5724 genes
ChIP-seq density Log2
ChIP-seq Log2 FC
RNA-seq Log2 FC
ChIP density
ChIP log2 FC
RNA log2 FC
H3K27ac
H3K4me1
RNA
Normal
Early 1
Early 2
Aggressive 1
Aggressive 2
Early 1 vs Normal
Early 2 vs Normal
Aggressive 2 vs Early 2
Aggressive 1 vs Early 1
Aggressive vs Early ex vivo
H
Enhancer overlap between stages
H3K27ac
Aggres-sive 2
Early 2
1,095
514
3,443
H3K4me1
2,781
730
6,997
I
Numbers of enhancers with early and late H3K27ac changes
1,936
2,021
148
190
160
16
Early changes
Later changes
J
Motifs enriched in differential enhancers
Spi1, IKZF1, ZFP14
Stat2, Bcl11B
PRDM9, ZNF148, ASCL1
ZNF384, ONECUT1
ELFs, EHF, Ikzf, ETV
NR5A1, NR5A2
KLF9
CEBPs
ELF5, GABPA, ELF3
FEV, ETS, ETV, GABPA
Runx1
JUN::JUNB
TFAP2A
Myb
KLF4, ZNF281, ZNF148
ZKSCAN3, ESR2
PITX1, PITX2
Aggr 1
Aggr 2
−10*log10(E-value)
0 200 400 600 800 1000

**Figure S3 (related to Figure 3). Changes of enhancer chromatin states at the aggressive stage were more clone-specific, skewed towards activation, and often reversed early leukemic changes. A-C.** Additional examples of ChIP-seq genomic tracks around enhancers that changed chromatin state at the aggressive stage. Input-normalized ChIP-seq densities of H3K27ac (green) and H3K4me1 (dark green) in normal progenitors (top) compared to two independent leukemic clones 1 and 2 at early and aggressive stages. Tracks are shown for 3 promoters Peg10 **(A)**, Pou4f1 **(B)**, Sod1 **(C)**. **D-F** Chromatin state progression in the space of H3K27ac vs H3K4me1 for enhancers shown in **A-C**. Points represent normal progenitors (grey) and early (blue) and aggressive (red) stages of two leukemic clones (1 and 2). The corresponding gene expression levels (RNA-seq RPKM) are shown as barplots. **G.** Expanded heatmap of changes (log2 fold change) at the aggressive stage for H3K27ac and H3K4me1 densities across all enhancers with a substantive level and > 2-fold change of at least one mark (middle), juxtaposed with the absolute levels of mark densities in normal progenitors and two leukemic clones at early and aggressive stage (left) and changes of proximal gene expression in vivo (log2 fold change of RPKM, right). Absolute levels and changes of chromatin marks at the aggressive stage were only partially consistent between two independent clones (two columns for each mark). **H.** Most enhancers affected at the aggressive stage underwent prior changes at the early stage. Data for clone 2. Venn diagrams of enhancers with changes of H3K27ac and H3K4me1 at the aggressive and early stages of the same clone. **I.** Clonal evolution preferentially reversed the prior early leukemic changes of H3K27ac at enhancers. Large arrows: numbers of enhancers with increase (red) and decrease (blue) of H3K27ac between normal progenitors and the early leukemic subclone. Smaller arrows near each large arrow: numbers of these enhancers that underwent the later increase (red) or decrease (blue) of H3K27ac during clonal evolution. These later changes were skewed in the direction opposite to the early leukemic changes. Data for clone 2. **J.** Enrichment of transcription factor binding motifs associated with enhancer mark changes during clonal evolution. MEME enrichment E-values for transcription factor motifs (rows) shown for enhancers changing the level of a given chromatin mark in a given clone (columns).

Figure S4

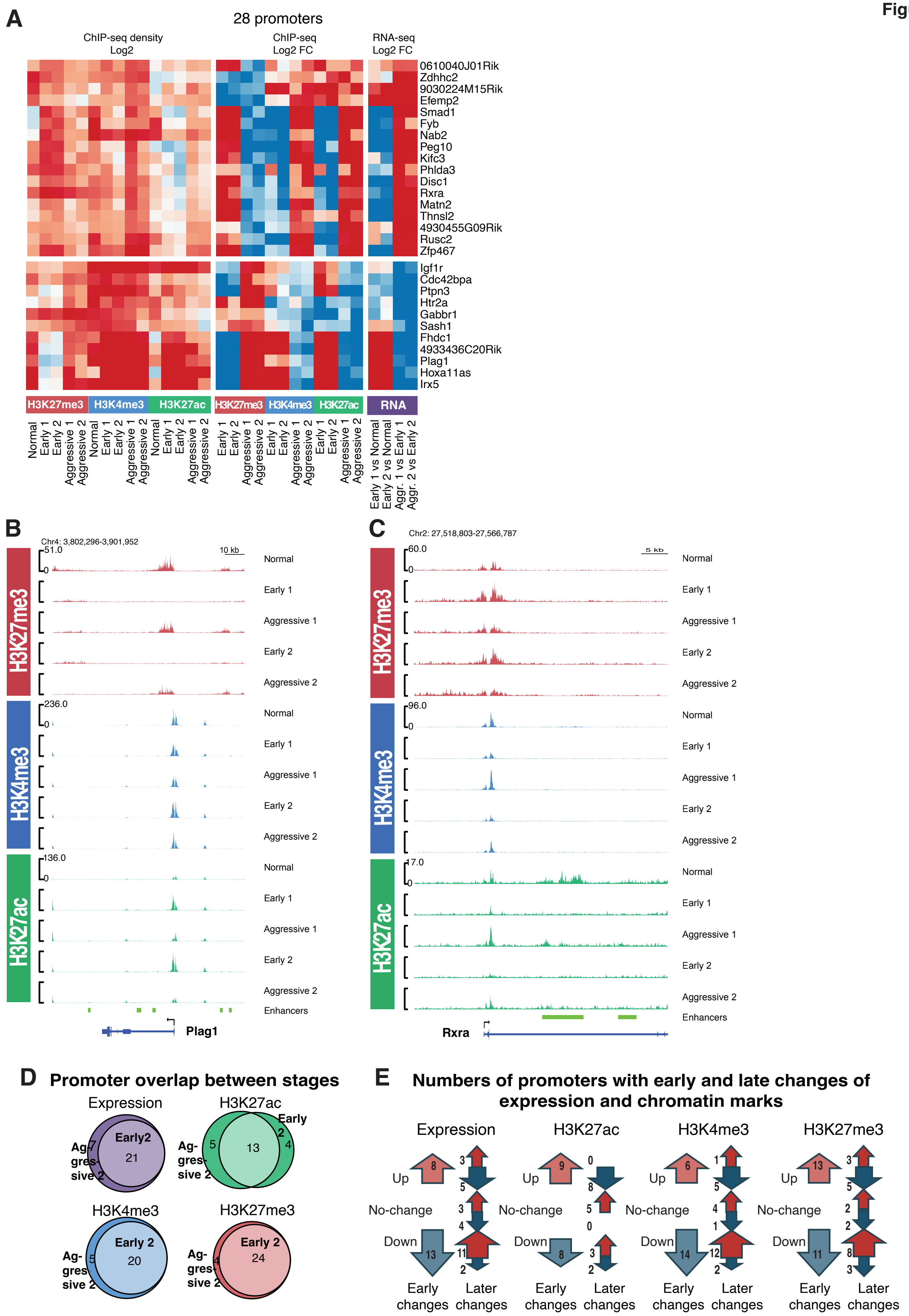
A
28 promoters
ChIP-seq density Log2
ChIP-seq Log2 FC
RNA-seq Log2 FC
0610040J01Rik
Zdhhc2
9030224M15Rik
Efemp2
Smad1
Fyb
Nab2
Peg10
Kifc3
Phlda3
Disc1
Rxra
Matn2
Thnsl2
4930455G09Rik
Rusc2
Zfp467
Igf1r
Cdc42bpa
Ptpn3
Htr2a
Gabbr1
Sash1
Fhdc1
4933436C20Rik
Plag1
Hoxa11as
Irx5
H3K27me3
H3K4me3
H3K27ac
RNA
Normal
Early 1
Early 2
Aggressive 1
Aggressive 2
Early 1 vs Normal
Early 2 vs Normal
Aggr. 1 vs Early 1
Aggr. 2 vs Early 2
B
Chr4: 3,802,296-3,901,952
10 kb
51.0
236.0
136.0
Enhancers
Plag1
C
Chr2: 27,518,803-27,566,787
5 kb
60.0
96.0
17.0
Rxra
D
Promoter overlap between stages
Expression
Early2
21
Ag-gres-sive 2
7
H3K27ac
Early 2
13
5
4
H3K4me3
Early 2
20
5
H3K27me3
Early 2
24
4
E
Numbers of promoters with early and late changes of expression and chromatin marks
Expression
H3K27ac
H3K4me3
H3K27me3
Up
No-change
Down
Early changes
Later changes

**Figure S4 (related to Figure 4). Changes of promoter chromatin state and expression among the core gene set at both stages of aggressiveness in two clones. A.** Expanded heatmap of promoter H3K27me3, H3K4me3, H3K27ac and expression changes between early and aggressive stages (fold change in log2 scale, right), juxtaposed with the changes at the same promoters between normal progenitors and early stage (left). **B-C.** ChIP-seq genomic tracks showing promoter chromatin mark and expression changes between two stages of aggressiveness for Plag1 and Rxra genes. H3K27me3 (red), H3K4me3 (blue), and H3K27ac (green) density tracks in normal progenitors (top) compared to two independent leukemic clones 1 and 2 at the early and aggressive leukemic stages. **D.** Virtually all core aggressiveness-associated genes underwent prior changes at the early leukemic stage. Data for clone 2. Venn diagrams of promoters with aggressiveness-associated (Aggressive) and early leukemic changes in the same clone (Early) for expression and individual chromatin mark. **E.** Clonal evolution preferentially reversed the prior early leukemic changes at the promoters of core genes. Large arrows: numbers of promoters with increase (red) and decrease (blue) of expression and chromatin marks (H3K27ac, H3K4me3, H3K27me3) between normal progenitors and the early leukemic subclone. Smaller arrows near each large arrow: numbers of these promoters that underwent the later increase (red) or decrease (blue) of expression or chromatin mark during clonal evolution. These later changes were skewed in the direction opposite to the early leukemic changes. Data for clone 2.

**Figure S5**

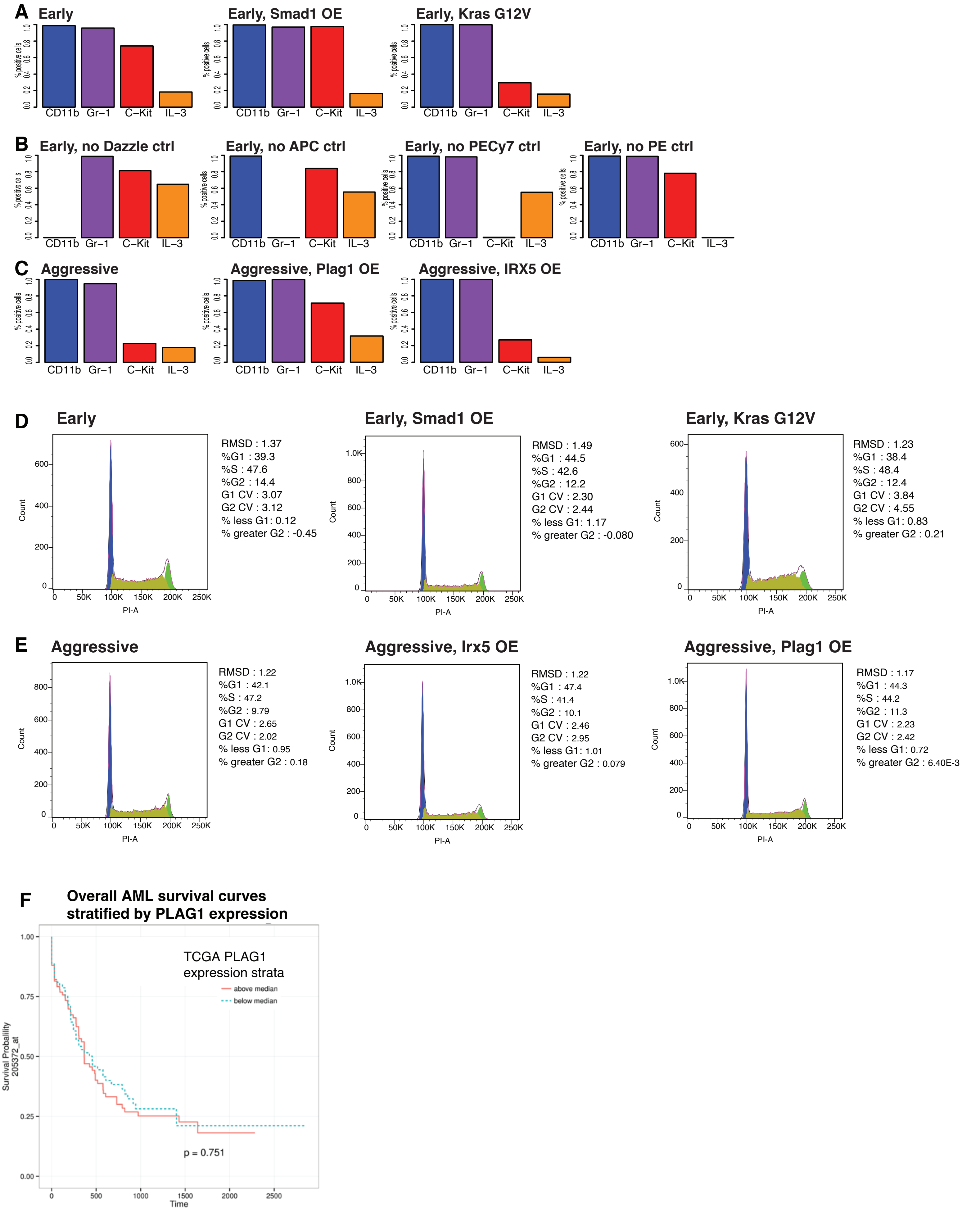
A
Early
Early, Smad1 OE
Early, Kras G12V
% positive cells
CD11b Gr−1 C−Kit IL−3
B
Early, no Dazzle ctrl
Early, no APC ctrl
Early, no PECy7 ctrl
Early, no PE ctrl
C
Aggressive
Aggressive, Plag1 OE
Aggressive, IRX5 OE
D
Early
RMSD : 1.37
%G1 : 39.3
%S : 47.6
%G2 : 14.4
G1 CV : 3.07
G2 CV : 3.12
% less G1: 0.12
% greater G2 : -0.45
Early, Smad1 OE
RMSD : 1.49
%G1 : 44.5
%S : 42.6
%G2 : 12.2
G1 CV : 2.30
G2 CV : 2.44
% less G1: 1.17
% greater G2 : -0.080
Early, Kras G12V
RMSD : 1.23
%G1 : 38.4
%S : 48.4
%G2 : 12.4
G1 CV : 3.84
G2 CV : 4.55
% less G1: 0.83
% greater G2 : 0.21
Count
PI-A
E
Aggressive
RMSD : 1.22
%G1 : 42.1
%S : 47.2
%G2 : 9.79
G1 CV : 2.65
G2 CV : 2.02
% less G1: 0.95
% greater G2 : 0.18
Aggressive, Irx5 OE
RMSD : 1.22
%G1 : 47.4
%S : 41.4
%G2 : 10.1
G1 CV : 2.46
G2 CV : 2.95
% less G1: 1.01
% greater G2 : 0.079
Aggressive, Plag1 OE
RMSD : 1.17
%G1 : 44.3
%S : 44.2
%G2 : 11.3
G1 CV : 2.23
G2 CV : 2.42
% less G1: 0.72
% greater G2 : 6.40E-3
F
Overall AML survival curves
stratified by PLAG1 expression
TCGA PLAG1
expression strata
above median
below median
p = 0.751
Survival Probability
205372_at
Time

**Figure S5 (related to Figure 5). A-B.** Flow cytometry analysis of lineage markers in Smad1, Irx5, and Plag1 transduced cells suggest no changes in hematopoietic lineage. Barplots showing fractions of Cd11b, Gr-1, c-Kit, and Il-3 positive cells among Early-stage cells. **A.** Overexpressing Smad1 and controls. **B.** Marker controls. **C.** Fractions among Aggressive-stage cells overexpressing Plag1 and Irx5, and controls. **D-E**. Cell cycle analysis from flow cytometry using propidium iodide DNA staining in transduced and non-transduced progenitors. Overexpression of Smad1, Irx5, and Plag1 resulted in only modest shifts between the phases of cell cycle. In the early subclone, Smad1 overexpression decreased S-phase fraction by 5%, from 48% to 43%, and increased G1 fraction by 5%, from 39% to 44%, suggesting a moderate extension of G1 at the expense of S-phase. In the aggressive subclone, both Irx5 and Plag1 overexpression produced similar small shifts towards the extension of G1-phase and decrease of S-phase. **F**. Analysis of public survival data among AML patients stratified by PLAG1 expression. Kaplan-Meier curves for AML patients with Plag1 expression levels above (red) and below median (blue). Analysis of TCGA AML patient cohort implemented in BloodSpot. Plag1 expression is not associated with strong survival differences in this patient cohort.
